# Genetically engineered mesenchymal stem cells as a nitric oxide reservoir for acute kidney injury therapy

**DOI:** 10.1101/2022.12.11.519950

**Authors:** Haoyan Huang, Meng Qian, Yue Liu, Shang Chen, Huifang Li, Zhibo Han, Zhong-Chao Han, Xiang-Mei Chen, Qiang Zhao, Zongjin Li

**Author notes:** Correspondence: Qiang Zhao, Nankai University, 94 Weijin Road, Tianjin 300071, China, or Zongjin Li, MD, Nankai University, 94 Weijin Road, Tianjin 300071, China. These authors contributed equally to this work (Haoyan Huang, Meng Qian).

## Abstract

Nitric oxide (NO), as a gaseous therapeutic agent, shows great potential for the treatment of many kinds of diseases. Although various NO delivery systems have emerged, the immunogenicity and long-term toxicity of artificial carriers hinder the potential clinical translation of this gas therapeutics. Mesenchymal stem cells (MSCs), with the capacities of self-renewal, differentiation, and low immunogenicity, have been used as living carriers. However, MSCs as gaseous signaling molecule (GSM) carriers have not been reported. In this study, MSCs were genetically modified to produce mutant β-galactosidase (β-GAL^H363A^). Furthermore, a new NO prodrug, 6-methyl-galactose-benzyl-oxy NONOate (MGP), was designed. MGP can entrance into cell and selectively trigger the NO release from genetically engineered MSCs (eMSCs) in the presence of β-GAL^H363A^. Moreover, our results revealed that eMSCs can release NO when MGP is systemically administered in a mouse model of acute kidney injury (AKI), which can achieve NO release in a precise spatiotemporal manner and augment the therapeutic efficiency of MSCs. This eMSC and NO prodrug system provides a unique and tunable platform for GSM delivery and holds promise for regenerative therapy by enhancing the therapeutic efficiency of stem cells.

**Significance:** Gaseous signaling molecules (GSMs), such as nitric oxide (NO), hold tremendous potential in regenerative medicine and tissue engineering. However, the delivery of GSMs with stem cells to target tissues or organs in response to demand has never been reported. This study designed engineered mesenchymal stem cells (eMSCs) that produce mutant β-galactosidase (β-GAL^H363A^) and trigger NO release when the NO prodrug is systemically administered, which can achieve NO release in a precise spatiotemporal manner and augment the therapeutic efficiency of MSCs in a mouse model of acute kidney injury (AKI). This eMSC and NO prodrug system provides a unique and tunable platform for GSM delivery and holds promise for regenerative therapy by enhancing the therapeutic efficiency of stem cells.

**Graphical abstract:** 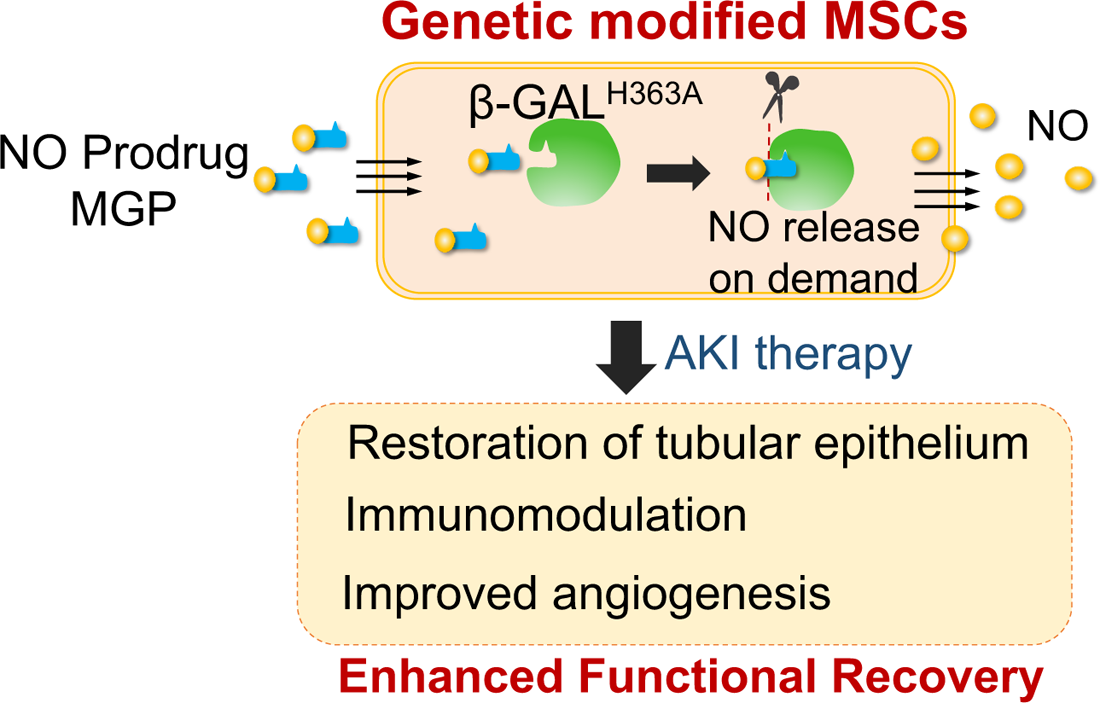

## INTRODUCTION

Nitric oxide (NO), as a gaseous signaling molecule, plays vital roles in various physiological processes, including tissue regeneration (1). Due to the extremely short half-life of NO, efforts to extensively enhance the therapeutic efficacy of NO by various types of artificial carriers, such as polymers, peptides, and nanoparticles, have achieved many successes, and there are remaining challenges that need to be addressed, including off-targeting, toxicity, immunogenicity, and clinical translation (2, 3). Successful NO delivery requires an appropriate carrier for delivering these therapeutic molecules to the target sites in an on-demand and controlled manner, as well as in a protected, pharmacologically active form(4). Bioinspired delivery vehicles, including biological cells, exosomes, and isolated membrane ghosts, are highly attractive because they exhibit excellent biodistribution, immune compatibility, innate disease-targeting abilities, and reduced toxicity(5–8). Among these systems, mesenchymal stem cells (MSCs) have gained considerable attention as drug-delivery vehicles owing to their genetic tractability, payload diversity, intrinsic tropism for disease sites, and differentiation potential (9, 10).

MSCs are adult stem cells capable of self-renewal and multilineage differentiation that have been reported to promote tissue regeneration mediated by immunomodulatory and proangiogenic properties via paracrine effects, and MSC-based cell therapeutics have entered multiple clinical trials(11). MSCs as delivery vehicles have been widely studied for the delivery of various drugs and bioactive molecules into target disease sites, including peptides, proteins, DNA, and RNA(12, 13). However, the concept of MSCs as vehicles to deliver gaseous signaling molecules (GSMs) has not been studied. In this regard, we propose an MSC-based gas-generating platform as a unique and tunable platform for extensively broad GSM therapies.

To achieve a gas-generating platform based on living cells, MSCs are programmed to sustain the generation and release NO by taking advantage of enzyme prodrug therapy (EPT), a versatile and exploitable technique to convert inactive, nontoxic prodrugs to active drugs at the desired sites(14). In the present study, we developed an engineered MSC-based NO delivery platform for controlled NO delivery. In this advanced delivery platform, the expression of mutant β-galactosidase (β-GAL^H363A^) by eMSCs enabled the production of NO when the NO prodrug was administered. We hypothesized that eMSCs could successfully generate and release NO in a precise spatiotemporal manner, as well as augment the therapeutic effects of MSCs and result in superior efficacy of stem cell therapy.

## RESULTS

### MSC-mediated NO release platform

To establish a living cell-mediated gas generation platform, MSCs were genetically modified to produce mutant β-galactosidase (β-GAL^H363A^), which triggers NO release from MSCs when the corresponding NO prodrug is applied and avoids the interference of endogenous glycosidase. We constructed a lentiviral vector to stably express mutant β-GAL^H363A^, Renilla luciferase (Rluc), and red fluorescence protein (RFP), in which Rluc and RFP were used for molecular imaging and immunohistology, respectively (**Fig. 1A**). The expression of β-GAL^H363A^ in engineered MSCs (eMSCs) was confirmed by Western blot analysis (**Fig. 1B**) and immunofluorescence analysis (**Fig. 1C**). Moreover, Rluc expression was confirmed by *in vitro* molecular imaging (**Fig. S1**). Furthermore, eMSCs showed similar morphology and expression patterns, indicating minimal side effects of genetic modification (**Fig. S2**).

**Fig. 1.**
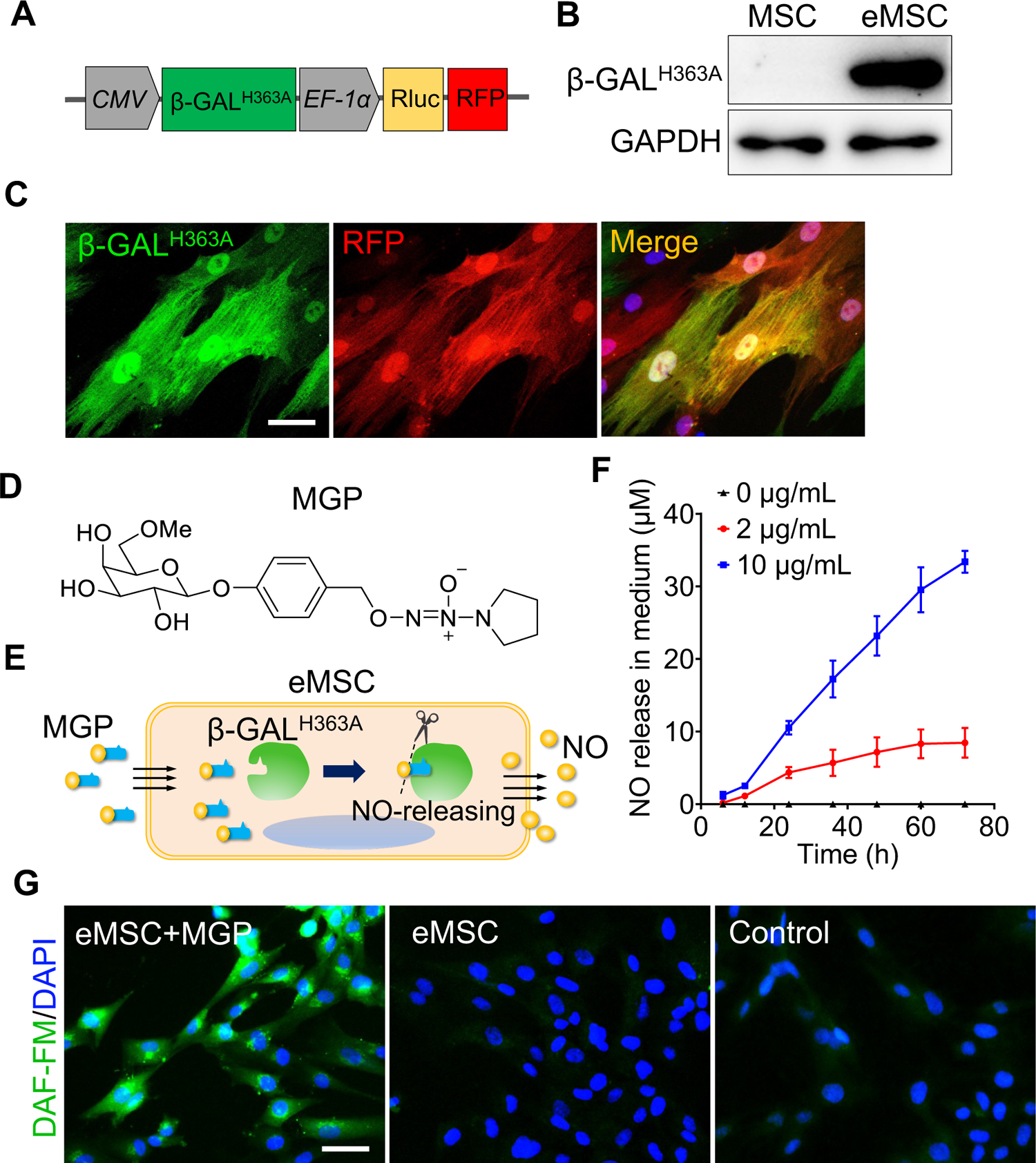
Evaluation of NO-eMSC system in NO generation and release. (**A**) Genetically modifying placenta-derived MSCs to express mutant β-galactosidase (β-GAL^H363A^) through a lentiviral transduction system and thus enabling continuous production of β-GAL^H363A^ for NO delivery. Moreover, the MSCs also stably expressed Rluc and RFP for *in vivo* bioluminescence imaging and immunofluorescence analysis, respectively. (**B**) The protein expression of β-GAL^H363A^ in eMSCs was evaluated by 6 × His tag protein detection due to β-GAL^H363A^ carrying a C-terminal 6×His tag. (**C**) Immunofluorescence staining of β-GAL^H363A^ (green) in eMSCs. Scale bars, 25 μm. (**D**) Structure of the NO prodrug MGP, 6-methyl-galactose-benzyl-oxy NONOate. (**E**) Construction of the NO-eMSC system and the mechanism of NO release from this system. (**F**) The NO release profile from eMSCs administered different concentrations of MGP was determined using the Griess assay. (**G**) The intracellular NO release in eMSCs was measured by DAF staining. eMSC+MGP, eMSCs with MGP administration. Scale bars, 50 μm.

Based on our previous report (15), we designed a new NO prodrug, 6-methyl-galactose-benzyloxy NONOate, which was named MGP. MGP can cross the cell membrane of eMSCs and then be catalyzed by β-GAL^H363A^ to release NO from eMSCs. The structure and synthesis routes of MGP are shown in **Fig. 1D** and **Fig. S3**. To verify whether the redesigned NO prodrug MGP could release NO specifically by the catalysis of β-GAL^H363A^, we measured the NO release profile of MGP in response to β-GAL^H363A^ *in vitro*. As shown in **Fig. S4**, MGP can be specifically catalyzed by β-GAL^H363A^ and has a sustained NO release behavior. In this novel NO release eMSC (NO-eMSC) system, eMSCs as a manufacturing factory for producing β-GAL^H363A^ could further release NO by catalyzing the NO prodrug MGP (**Fig. 1E**). We detected NO production in the medium of eMSCs with MGP administration using the Griess method, confirming that MGP can diffuse into eMSCs and release NO (**Fig. 1F**). Moreover, intracellular NO production was visualized by the fluorescent probe diaminofluorescein (DAF)- FM diacetate when eMSCs were treated with MGP, which also supported that eMSCs could specifically release NO (**Fig. 1G**).

Considering the cell-protective effects of a low NO concentration, our results revealed that the optimum concentration of MGP was 2 µg/mL by cell viability, immunostaining for the proliferation marker Ki67, and immunostaining for the expression of proliferation-related genes (**Fig. S5**). We further verified the protective effects of NO in response to oxidative stress-induced eMSC apoptosis. The bioluminescence imaging (BLI) assay showed that eMSC proliferation was markedly ameliorated H_2_O_2_-induced oxidative stress (**Figs. S6**), which was further confirmed by Ki67 staining and apoptotic analysis (**Figs. S7**).

### Enhanced antioxidation properties of eMSCs

To further investigate the transcriptomic profile of eMSCs with MGP administration, mRNA sequencing analysis was performed. We identified a total of 228 differentially expressed genes (DEGs) of which 116 were upregulated genes and 112 were downregulated genes (**Fig. S8A**). Analysis of gene ontology (GO) categories was mainly enriched in biological processes, including protein transport, cell differentiation, morphogenesis, and oxidation-reduction processes (**Fig. 2A**). Among them, the oxidation-reduction process attracted our attention, suggesting that NO may improve intracellular antioxidant capacity, which was confirmed by Gene Set Enrichment Analysis (GSEA) (**Figs. 2B and S8B**). Meanwhile, we noted that a significant increase in some of the DEGs associated with antioxidation was observed in eMSCs with MGP administration, namely, GSR, SRXN1, RRBP1, BMI1, RPL24P4, and MAFG, indicating that NO may indeed play a cytoprotective role in eMSCs by elevating antioxidation capacity (**Fig. 2C**). We also found that NO led to robust increases in survival- or proliferation-related genes, including FER, NRSN2, UCK2, CAB39, SNHG4, and SYDE1 (**Fig. 2D**), as well as angiogenesis-related genes, such as ARRB2, RHOQ, MYADM, and CFHR1 (**Fig. S8C**). Consistent with the aforementioned findings, GSEA of the whole transcriptome also revealed negative regulation of cell apoptosis and positive regulation of angiogenesis by NO (**Figs. 2E and S8D**). In addition, KEGG analysis highlighted pathways that may contribute to enhanced antioxidant capacity, including the Jak-STAT signaling pathway and MAPK signaling pathway (**Fig. S8E**). Together, these results demonstrated that NO might be able to ameliorate the oxidative stress of eMSCs, thus elongating the survival of eMSCs (**Fig. 2F**).

**Fig. 2.**
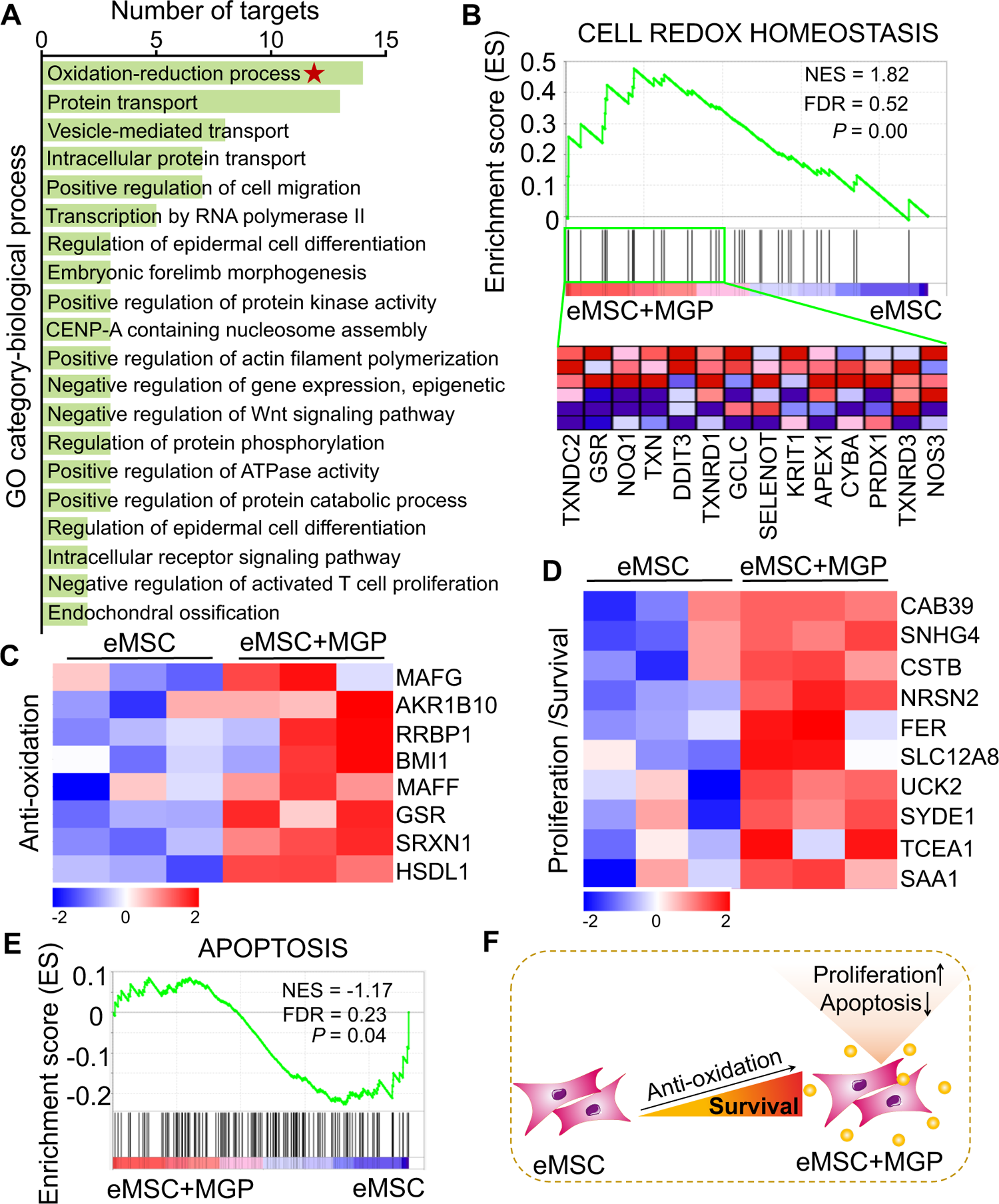
RNA-seq analysis revealed that NO improves the antioxidant capacity of eMSCs. (A) Gene Ontology (GO) category analysis of DEGs for biological processes. (**B**) Gene set expression analysis (GSEA) revealed enrichment for the cell redox homeostasis pathway. Heatmap of representative antioxidation (**C**) and proliferation/survival (**D**)-related genes. (**E**) GSEA revealed enrichment for apoptosis pathways. (**F**) Schematic representation of the prosurvival potential of the NO-eMSC system. eMSC+MGP, eMSCs with MGP administration; FDR, false discovery rate; NES, normalized enrichment score.

### Enhanced renoprotection of eMSCs

To further validate whether NO could promote eMSC survival *in vivo,* BLI analysis was performed for the real-time longitudinal monitoring of eMSC survival. A robust BLI signal of eMSCs was observed, indicating successful eMSC transplantation (**Fig. 3A**). Although all groups experienced donor cell death gradually in the following days, eMSCs with MGP administration strikingly exhibited increased cell retention and prolonged cell survival in comparison with the eMSC group (**Fig. 3B**). Furthermore, immunohistology also confirmed that the eMSCs with MGP administration displayed higher cell proliferation and cell retention at day 3 (**Fig. 3C**). Moreover, we found that the eMSCs treated with MGP showed significantly lower levels of SCr and BUN (**Figs. 3D and E**). Further kidney histology examination showed expanded proximal tubules, cast formation, and massive loss of brush borders at the initial stage after injury (3d) (**Figs. 3F to H**). Kidney fibrosis analysis demonstrated that the development of interstitial fibrosis was consistent with the mRNA expression of ECM synthesis- and fibrosis pathway-related genes (**Fig S9**).

**Fig. 3.**
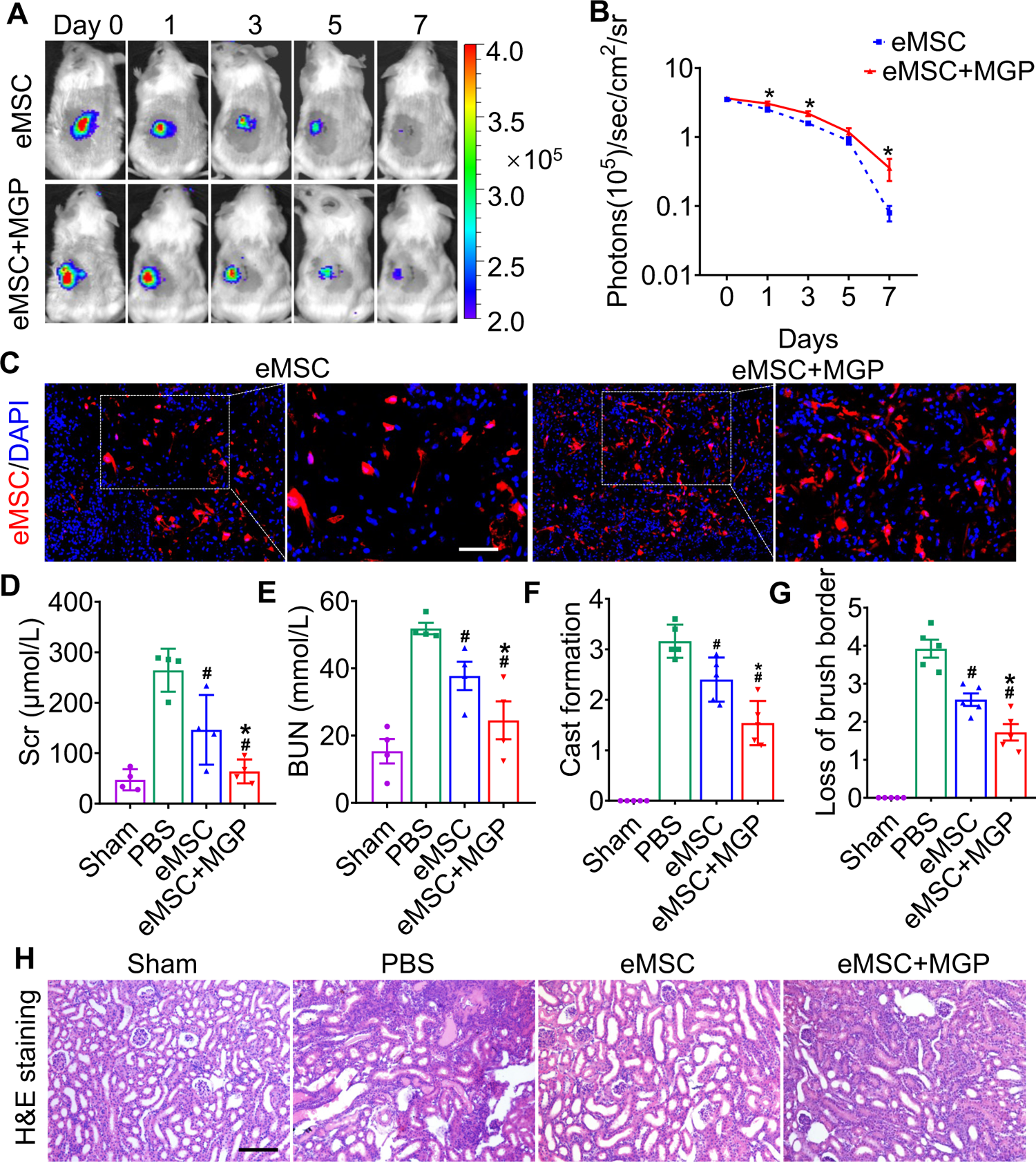
The NO-eMSC system confers renoprotection by increasing eMSC survival *in vivo*. (**A and B**) The survival ratio of eMSCs and eMSCs with MGP administration after transplantation in AKI model mice was tracked by BLI. The number of transplanted eMSCs was 1 × 10^6^ cells. (**C**) Representative photomicrographs show the retention ratio of transplanted eMSCs (red) and NO-eMSCs (red) on day 3 post-AKI. Cell nuclei were stained with DAPI (blue). Scale bar, 100 μm. Renal function indexes, including serum creatine (**D**) and blood urea nitrogen (**E**) levels, were measured on day 3 after AKI. (**C**) Quantitative analysis of cast formation (**F**) and loss of brush border (**G**) of H&E staining. (**H**) Representative images of H&E staining on day 3 post AKI. eMSC+MGP, eMSCs with MGP administration. Scale bar, 100 μm. All data are presented as the mean ± SD, n≥3 per group, \**P* < 0.05.

Furthermore, the expression of KIM-1, a kidney injury marker, was significantly decreased in the proximal tubules of the kidneys in the eMSCs with MGP administration group compared to the eMSC group (**Figs. S10A and B**). Simultaneously, we found that the level of caspase-3/cleaved caspase-3 exhibited an abrupt increase in the injured kidney of the PBS group but was attenuated by eMSCs with MGP administration (**Figs. S10C and D**), suggesting that the NO-eMSC system has protective effects on apoptosis in tubular epithelial cells. Overall, these results indicated that the NO-eMSC system could increase eMSC survival in vivo and further improve kidney function.

### Evaluation of NO levels in kidney

To further evaluate *in vivo* NO release from the eMSC, eMSCs were transplanted into the left kidney followed by intravenous injection of the NO prodrug MGP. The results from Griess (**Fig. 4A**) and chemiluminescence assays (**Fig. 4B and C**) showed that the NO level from eMSC-treated kidneys was significantly increased when the prodrug MGP was applied. We next directly measured the NO levels of eMSC-treated kidneys *in vivo* by an EPR technique using the spin trap ferrous N-methyl-d-glucamine dithiocarbamate complex ((MGD)_2_Fe^2+^). The EPR signal of NO was observed in all groups, while an obvious characteristic triplet EPR signal (NO signal) was observed in the eMSCs treated with MGP (**Fig. 4D and E**), which further confirmed that eMSCs could release NO *in vivo*. Moreover, targeted NO delivery was evaluated by comparison of NO levels in different tissues of mice after the administration of the eMSC in the kidney. The quantitative data showed that NO production from the kidney was significantly higher than that from the heart and liver (**Fig. 4F to H**), suggesting that eMSCs could release NO specifically in the kidney.

**Fig. 4.**
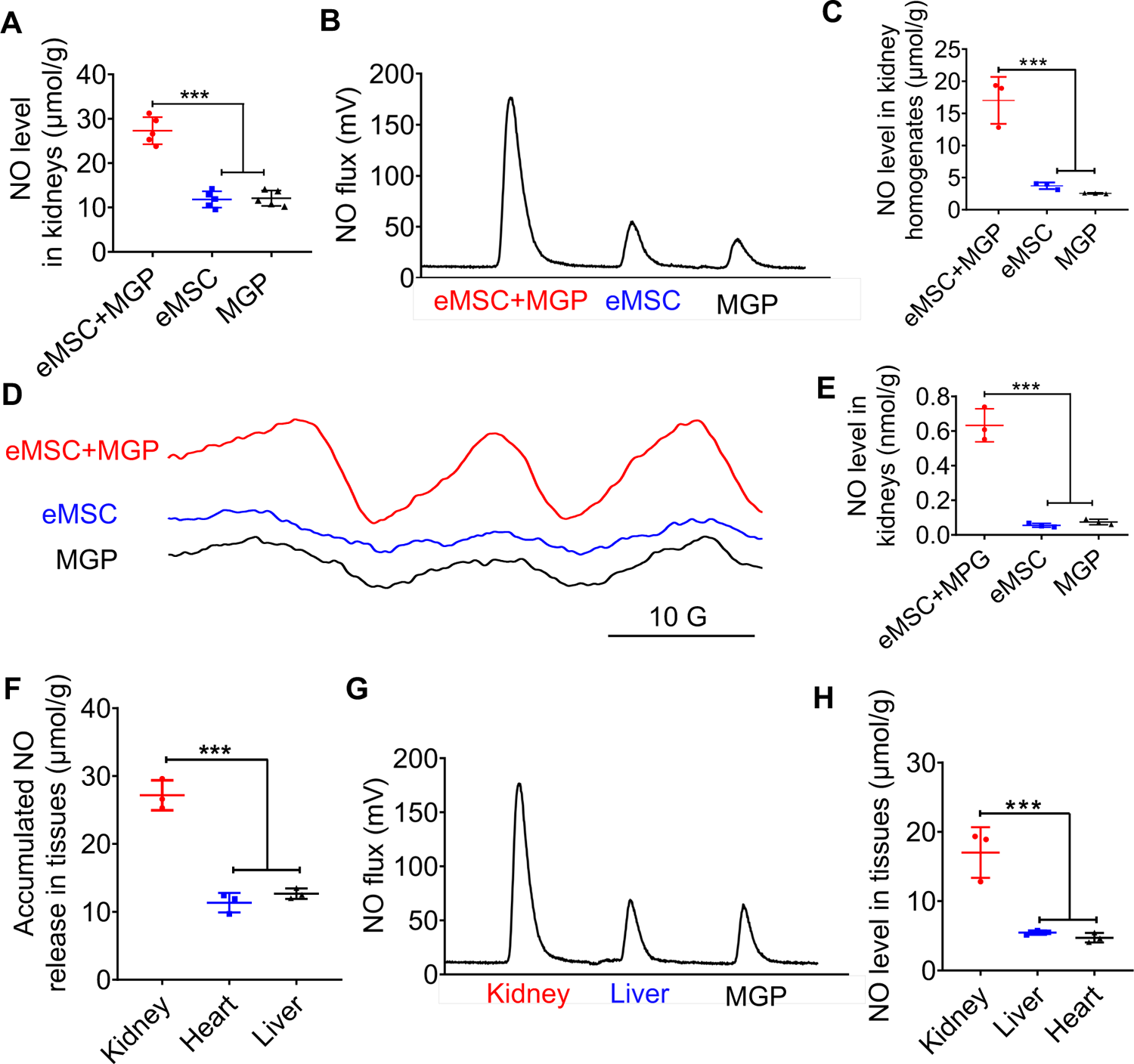
Detection of NO release from the NO-eMSC system *in vivo*. (**A**) The Griess reagent system was used to measure NO levels in kidneys. (**B** and **C**) Detection and quantification of NO levels in kidneys *in vivo* using chemiluminescence. (**D** and **E**) NO release from eMSCs in kidneys was assessed by EPR. (**F**) Griess reagent was used to measure NO levels in different tissues. Detection (**G**) and quantification (**H**) of NO levels in different tissues from NO-eMSC system-treated mice *in vivo* using chemiluminescence. eMSC+MGP, eMSCs with MGP administration. All data are presented as the mean ± SD, \**P* < 0.05, n≥3 per group.

### Enhanced proangiogenic effects of eMSCs

The VEGFR2-Fluc transgenic mice were used to monitor renal angiogenesis in real time, which appeared as BLI signals. According to the results of angiogenesis imaging, BLI signals were emitted in all groups, and the strongest signal was detected in the eMSCs with MGP administration group, which suggests that the NO-eMSC system could stimulate renal angiogenesis by activating the VEGF/VEGFR2 pathway (**Figs. 5A and B**). We next verified neovascularization in damaged kidneys on day 7 post-surgery by histologic examination. The CD31 immunostaining results revealed significantly enhanced CD31^+^ microvascular density in the group of eMSCs with MGP administration, which is consistent with the BLI results (**Figs. 5C and D**). Moreover, the expression of the angiogenesis-related genes *VEGFR2, bFGF, PLGF, Ang-1,* and *Ang-2* detected by real-time RT‒PCR also confirmed that the eMSCs with MGP administration facilitated renal angiogenesis by upregulating angiogenic factor expression (**Fig. S11**). Recovery from renal ischemia reperfusion (I/R) injury requires tubular cell proliferation to promote kidney regeneration. As shown in **Figs. 5E and F**, the numbers of proliferating Ki-67+ tubular cells were largely stimulated in the group of eMSCs treated with MGP, suggesting that the NO-eMSC system promoted kidney regeneration by accelerating tubular cell proliferation. The promotion of renal angiogenesis contributes to the restoration and repair of damaged kidneys, which paves the way for successful kidney regeneration.

**Fig. 5.**
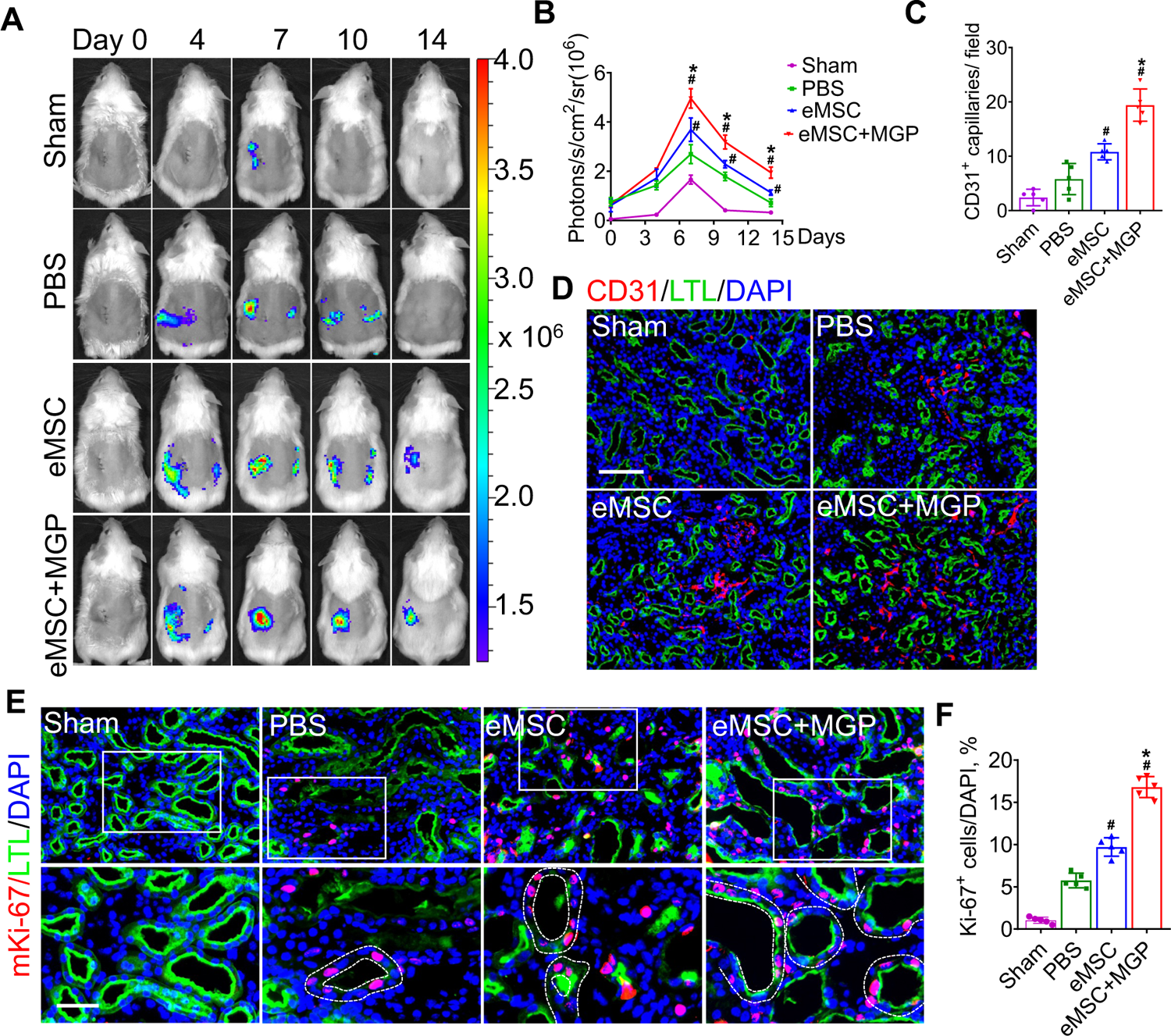
The NO-eMSC system promoted kidney regeneration after AKI. (**A**) BLI monitored the spatiotemporal dynamics of VEGFR2 expression following eMSC administration in a mouse AKI model. (**B**) Quantification of the BLI signals showed better angiogenic effects in the eMSCs with MGP administration. (**C**) Quantification of CD31-positive capillaries on day 7 after AKI. (**D**) CD31 staining was performed on day 7 after AKI. Scale bar, 100 μm. LTL (green) was used to reveal proximal tubules. Representative images of Ki67 staining (**E**) and percentage of Ki67-positive cells (**F**) on day 3 after AKI. eMSC+MGP, eMSCs with MGP administration. Scale bar, 50 μm. All data are presented as the mean ± SD, n≥3 per group, \**P* < 0.05 versus eMSCs; ^#^*P* < 0.05 versus PBS.

Subsequently, we explored the mechanism of eMSCs with MGP administration-induced angiogenesis, and our data revealed that eMSCs boosted the migration ability of HUVECs in a co-culture system with Transwell, as evidenced by a scratch wound healing assay (**Fig. S12A and B**). Additionally, the angiogenic ability of HUVECs, as manifested by the number of nodes and branches, was visibly increased in the group of eMSCs treated with MGP, as assessed by a tube formation assay (**Fig. S12C, D, and E**). Additionally, the protein level of cleaved caspase-3, a key apoptosis molecule, was detected by immunofluorescence staining, suggesting that eMSCs could inhibit the apoptosis of HUVECs (**Fig. S12F**). In general, we proposed that eMSCs with MGP administration exert superior promotive effects on kidney angiogenesis after injury by enhancing the proangiogenic activities of vascular endothelial cells.

### Diminished modulated inflammatory responses

We next explored the modulatory effect of eMSCs on the inflammatory response in I/R-injured kidneys, and our results showed that eMSCs with MGP administration increased the number of F4/80- and CD206-positive macrophages while decreasing the number of F4/80- and iNOS-positive macrophages, indicating that the NO-eMSC system could promote the polarization of M1 macrophages to M2 macrophages (**Fig. 6A and B**). Meanwhile, mRNA expression profiling further confirmed this finding (**Fig. 6C and D**). Oxidative stress-induced reactive oxygen species (ROS) are well known as the main cause of renal I/R injury, leading to subsequent augmented inflammation and extended tissue damage(16). Damaged renal tubular cells were overloaded with excessive lipids caused by abnormal lipid metabolism, while elevated ROS levels can lead to irreversible oxidative damage to lipids and further result in cell damage and death. Excessive intracellular accumulation of lipids was observed in the renal tubules of the PBS group after I/R injury, while the eMSCs with MGP administration significantly decreased lipid deposition **(Fig. S13A and B)**. Meanwhile, the renal level of MDA, an index of ROS-mediated lipid peroxidation, was notably increased in the PBS group and significantly decreased in the eMSCs with MGP administration group (**Fig. S13C**). Moreover, we examined the antioxidant capacity levels (SOD and GSH contents) in kidney tissues, and remarkable upregulation of SOD and GSH levels was observed in the group of eMSCs treated with MGP (**Fig. S13D and E**). In conclusion, the NO-eMSC system exhibited a renal protective effect by targeting antioxidation and macrophage polarization-mediated anti-inflammation.

**Fig. 6.**
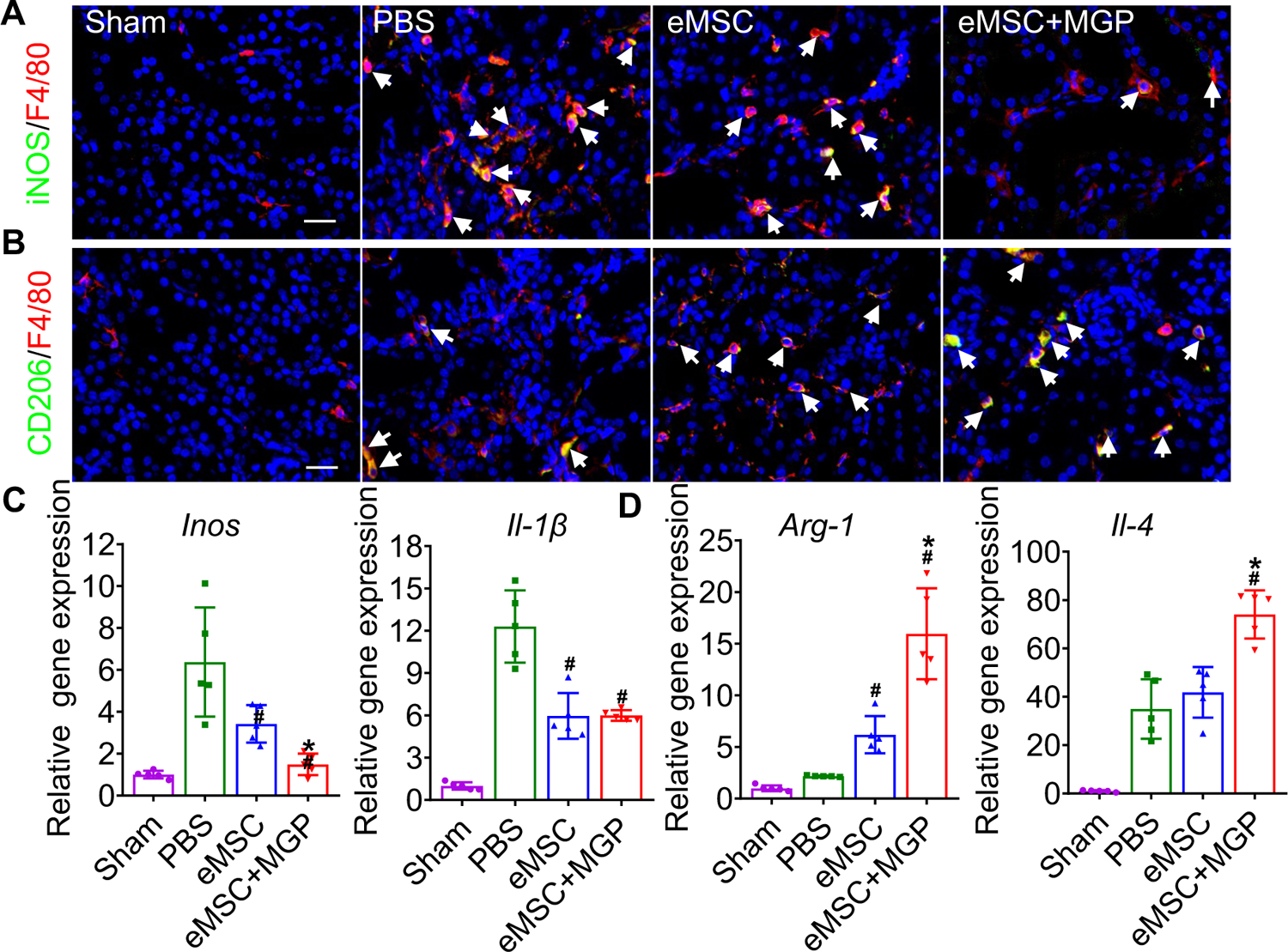
The NO-eMSC system modulates the inflammatory response by reshaping macrophage polarization. (**A**) Representative images of immunostaining for the M1 macrophage markers iNOS (green) and F4/80 (red) on day 3 after AKI. Scale bar, 100 μm. (**B**) Representative images of immunostaining for the M2 macrophage markers CD206 (green) and F4/80 (red) on day 3 after AKI. Scale bar, 100 μm. (**C**) RT‒PCR analysis of M1-related gene (iNOS and IL-1β) expression in kidney tissues at day 3 after AKI. (**D**) RT‒PCR analysis of M2-related gene (IL-4 and Arg-1) expression in kidney tissues at day 3 after AKI. Scale bar, 100 μm. eMSC+MGP, eMSCs with MGP administration. All data are presented as the mean ± SD, n≥3 per group, \**P* < 0.05 versus eMSCs; ^#^*P* < 0.05 versus PBS.

## DISCUSSION

In this study, we have developed an eMSC-mediated NO gas-generating platform, the NO-eMSC system, for site specific, controlled, and long-term delivery of NO for the first time. This advanced NO-eMSC system was engineered to both program the continuous release of NO and enhance stem cell-mediated regeneration of damaged tissues, which is a unique and mosaic approach to maximize the therapeutic effect. The NO-eMSC system successfully generated and released NO in a precise spatiotemporal manner and improved eMSC functionalities, resulting in higher cell survival and more resistance to apoptosis under oxidative stress. The NO-eMSC system exerted superior therapeutic effects, including rescuing renal tubules, relieving inflammation, stimulating angiogenesis, and improving renal function.

Gaseous signaling molecules (GSMs) with pharmaceutical effects, such as oxygen, NO, carbon monoxide, hydrogen, and hydrogen sulfide (H_2_S), have tremendous potential in the treatment of many diseases(17–19). Notably, the therapeutic effects of GSMs are highly concentration dependent; for example, low levels of NO (nmol to μmol) are frequently associated with cell protection, while high concentrations (>mmol) of NO exert apoptotic effects(1). Although a plethora of GSM small molecule donors has been developed, the insufficient solubility, poor stability, and unfavorable pharmacokinetics of these GSM donors have prevented their wide application(18). NO, the first and most well-known gaseous molecule, has been widely shown to play critical roles in a wide spectrum of physiological and pathological processes(18, 20). As a result, many kinds of NO-releasing platforms have emerged in the field of chemical materials to deliver NO in a controlled manner(18, 21). Although endogenous (enzyme and pH variations) or exogenous (light and ultrasound)-triggered NO delivery systems for controlled NO release profiles have been developed, it remains a challenge to maintain long-term NO release and eliminate immunogenicity and systemic toxicity(21, 22).

In recent years, an increasing number of cell delivery systems have entered multiple clinical trials or been approved for marketing(23, 24). Genetic engineering of MSCs is an approach to produce and deliver specific therapeutic proteins or enhance innate properties such as migration, differentiation, and survival for treating many tremendous diseases(25). Therefore, genetically engineered MSCs, usually as living “drug reservoirs”, offer long-term and stable production of the therapeutic protein of interest and thus have gained appreciable attention as therapeutic delivery vehicles(5). Taking advantage of this NO delivery system and stem cell therapy, we engineered MSCs as an enzyme reservoir that catalyzes the NO-prodrug to produce NO on demand, which combines the therapeutic potential of MSCs and NO.

For NO delivery, the undesired release of NO was observed due to the widespread distribution of endogenous β-GAL in blood and other tissues(15). Our previous study developed a novel NO-enzyme-prodrug pair for the specific release of NO, in which native β-GAL is transformed into mutant homologs with a “hole” and only interacts with its corresponding “bump”-prodrug(15). Hence, genetically engineered MSCs as living reservoirs enabled the continuous production of the “hole”-enzyme (β-GAL^H363A^), thereby achieving the controlled release of NO from its corresponding NO-prodrug, MGP. Moreover, the NO detection results confirmed that our NO-generating platform could release NO in a precise spatiotemporal manner; the duration and dosing of NO release could be tuned by controlling the NO prodrug. Specifically, NO release is initiated in the presence of the NO prodrug as long as eMSCs survive. NO dosing is controlled by the concentration of the administered NO prodrug, and an elevated NO signal was observed only at the site of transplanted eMSCs. In this study, we substantially advance the concept of a cell-mediated gas-generating platform for NO long-term applications, which may be applied for other gas enzyme prodrug pairs in the future.

The combinational use of NO-releasing biomaterials and stem cells is an emerging field of regenerative medicine(20); however, temporal and local release of NO-releasing biomaterials results in less contact time with cells and hinders their long-term applications. Moreover, in-depth knowledge of the underlying mechanisms to elucidate NO influence on stem cells and the stem cell microenvironment is of great value in the fields of biomaterials and regenerative medicine(1, 20). In this study, we developed an advanced MSC-mediated NO-generating platform engineered to both program the continuous release of NO and enhance stem cell-mediated regeneration of damaged tissues. Moreover, the NO-eMSC system exerted an immunosuppressive effect by reshaping macrophage inflammatory properties and an enhanced angiogenesis effect to promote kidney regeneration. This endogenous-like NO-generating platform may represent a new paradigm for the future of long-term NO application on MSCs or other cell types.

In conclusion, we have developed a genetic eMSC-based NO delivery, NO-eMSC system, for targeted, controlled, and long-term NO delivery for the first time. This advanced NO delivery system can serve as a generic delivery system that holds promise for extensively broad other gaseous molecule therapies and provides a new library of clinical solutions for regenerative medicine.

## Materials and Methods

Detailed Materials and Methods are provided in the SI Appendix. The synthesis of the MGP-NO prodrug is described in **Supplementary Note 1**.

### Cell culture

The MSCs used in this study were isolated from human placental tissue as previously described (26). The MSCs were transduced with lentiviral vectors carrying β-GAL^H363A^(15), Rluc, and RFP. Stable cells were isolated using FACS for RFP expression. The Rluc activity of engineered MSCs (eMSCs) was confirmed *in vivo* and *in vitro* using the IVIS Lumina II system (Xenogen Corporation, Hopkinton, MA). The cell surface marker expression of eMSCs was tested using a FACSCalibur flow cytometer (BD Biosciences).

### NO release detection in eMSCs

For *in situ* visualizations of intracellular NO, a specific membrane-permeable NO molecular probe, DAF-FM diacetate, was used, which was loaded into cells and produced DAF-FM by intracellular esterase(15). Cultured eMSCs were incubated with 5 μM DAF-FM diacetate at 37 °C for 20 min followed by extensive washing and then incubated with 5 μg/mL NO prodrug at 37 °C for 6 h, according to the manufacturer’s instructions (S0019; Beyotime).

### Assessment of NO release *in vivo* by chemiluminescence

Chemiluminescence methods were adopted to measure NO release from tissues by using a nitric oxide analyzer (NOA 280i, Zysense, USA). On day 7 after injury, the mice were sacrificed, and the samples were harvested. Then, the tissues were homogenized, and the supernatant was collected for NO detection. The samples were immersed in 5 mL of vanadium (III) chloride (50 mM), and the generated NO gas was diffused in the test solution and transported to the NO analyzer by a stream of N_2_. The generated NO was calculated using NaNO_3_ as a standard.

### NO release in the supernatant of cell culture

NO production in culture media and tissues was measured using the Nitric Oxide Assay Kit (S0021; Beyotime) based on the Griess reaction according to the manufacturer’s instructions. For *in vitro* NO release, 5×10^3^ eMSCs were seeded in 24-well plates and then incubated with different concentrations (0, 2, 10 μg/mL, 1 mL/per well) of the NO-prodrug MGP. At each predetermined time interval, 50 μL of the culture media was transferred to a 96-well plate, and 50 μL of Griess I and 50 μL of Griess II were added in sequence. Finally, a microplate reader (Promega) was used to measure the optical density (OD) values of the samples at 540 nm.

### Measurement of NO release *in vivo* by the EPR technique

The NO production of tissues *in vivo* was measured by electron paramagnetic resonance (EPR) spectroscopy as previously described(27). C57BL/6 albino mice (20-25 g) were anesthetized using 1×10^6^ cells (30 μL) of eMSCs transplanted into the left kidney through renal parenchymal injection. Then, DETC sodium salt (500 mg/kg, Sigma‒Aldrich) dissolved in water (250 mM) was injected subcutaneously. After 5 min, ammonium ferrous sulfate (50 mM) dissolved in citrate solution (250 mM) was administered by subcutaneous injection (2 mL/kg). Subsequently, 200 μL of 1 mg/mL MGP was administered via tail vein injection and allowed to circulate for 1 hour. Tissues were harvested and quickly frozen in liquid nitrogen. After that, the samples were crumbled into small pieces and extracted with 200 μL of ethyl acetate immediately. X-band EPR was used to measure the ethyl acetate extract in 50 μL capillary tubes at room temperature. Instrument settings were as follows: modulation frequency, 100 kHz; microwave power, 10 mW; modulation amplitude, 2 G; the number of scans, 30. The double integrated area of the EPR spectrum was calibrated into concentrations of NO-Fe^2+^(DETC)_2_ using TEMPO as a standard.

### RNA sequencing

RNA-seq samples of eMSCs and eMSCs treated with MGP (2 μg/mL, 24 h) were prepared and sent to LC Bio Tech (China) for analysis. Differentially expressed genes (DEGs) among the different samples were defined by the DESeq2 package as criteria of fold change (FC) ≥ 1.5 or FC ≤ 0.67 and P value ≤ 0.05. Functional enrichment analysis of GO and KEGG were based on the Gene Ontology Database (http://www.geneontology.org/) and KEGG pathway database (http://www.genome.jp/kegg/), respectively. GSEA/MSigDB databases were used for the GSEA (Broad Institute, http://www.broadinstitute.org/gsea/msigdb/index.jsp).

### Mice and Animal model

C57BL/6 albino mice (10 weeks old, female) were purchased from the Charles River Laboratories (Charles River). The I/R injury-induced AKI model was established as previously described(28). After reperfusion, 1×10^6^ cells (30 μL) of eMSCs were transplanted into the left kidney through renal parenchymal injection. Subsequently, 200 μL of 1 mg/mL NO prodrug, MGP, was administered via tail vein injection and then once for the other day until the end of the experiment. To monitor angiogenesis in real time *in vivo*, a C57BL/6 albino and outbred (Nu/Nu) background Vegfr2-luciferase transgenic mice (10 weeks old, female) were used to establish an AKI model induced by I/R injury (29). This study was performed in strict accordance with the recommendations in the Guide for the Care and Use of Laboratory Animals of the National Institutes of Health. All of the animals were handled according to the Nankai University Animal Care and Use Committee Guidelines (approval no. 2021-SYDWLL-000426).

### Statistical analysis

To check for significant differences, one-way analysis of variance (ANOVA) was used followed by Tukey’s HSD test when needed to compare three or more experimental groups of data, and Student’s t test was used to compare two groups of data. All data were analyzed using SPSS Statistics, version 25.0 (IBM SPSS Inc., Chicago, IL, USA). All quantification data are presented as the mean ± SD of at least three independent replicates.

### Data and Materials Availability

Supplementary study data are online and materials used in this study are available from the corresponding author upon reasonable request. All study data are included in the article and/or supporting information. Sequence data is available in the NCBI Sequence Read Archive under the primary accession code PRJNA910491 (https://www.ncbi.nlm.nih.gov/bioproject).

## ACKNOWLEDGMENTS

This study was financially supported by the National Key R&D Program of China (2017YFA0103200), the National Natural Science Foundation of China (U2004126), the Tianjin Natural Science Foundation (21JCZDJC00070), and Tianjin Key Medical Discipline (Specialty) Construction Project (TJYXZDXK-043A).

## Appendix 1--Key resources table

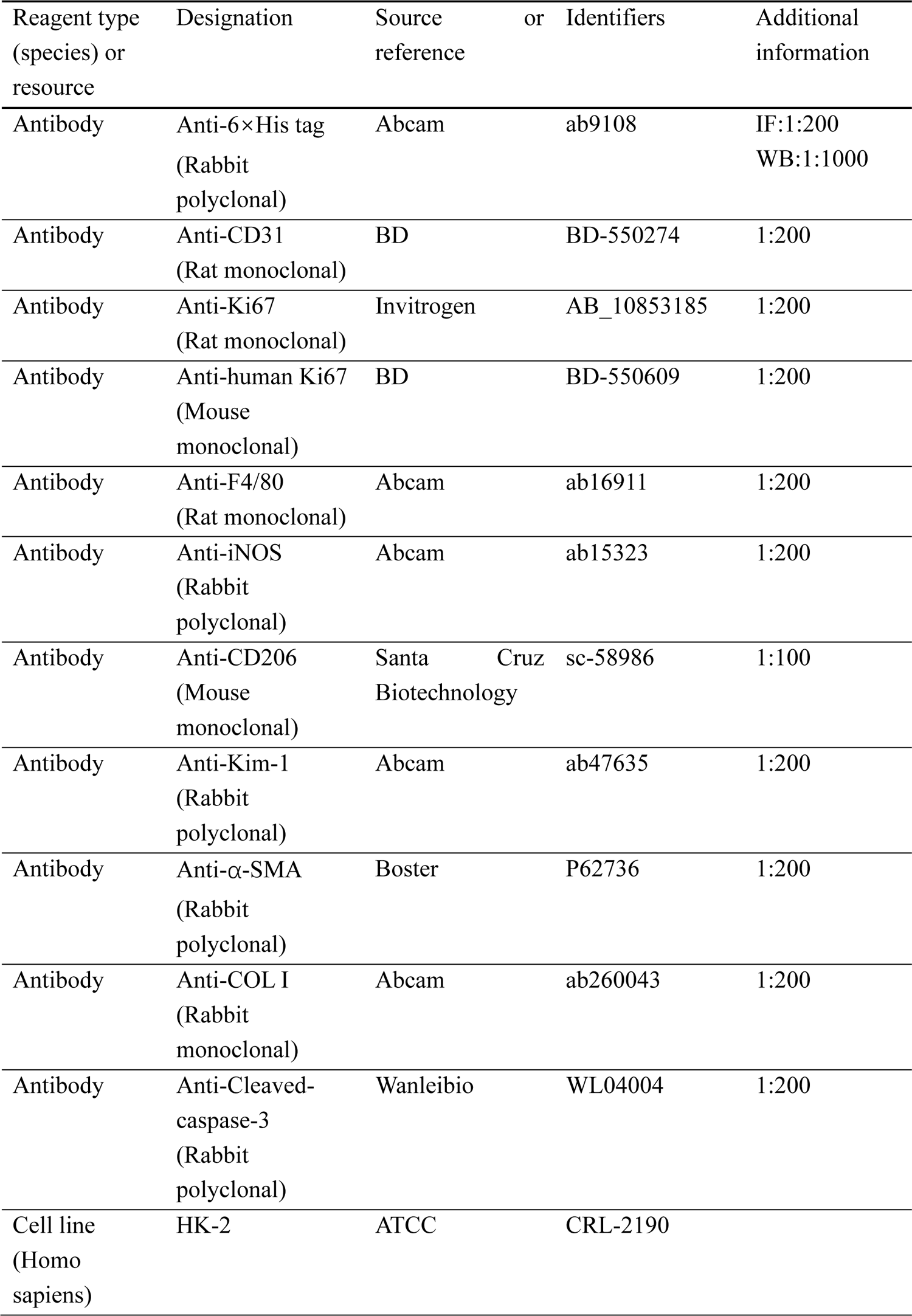
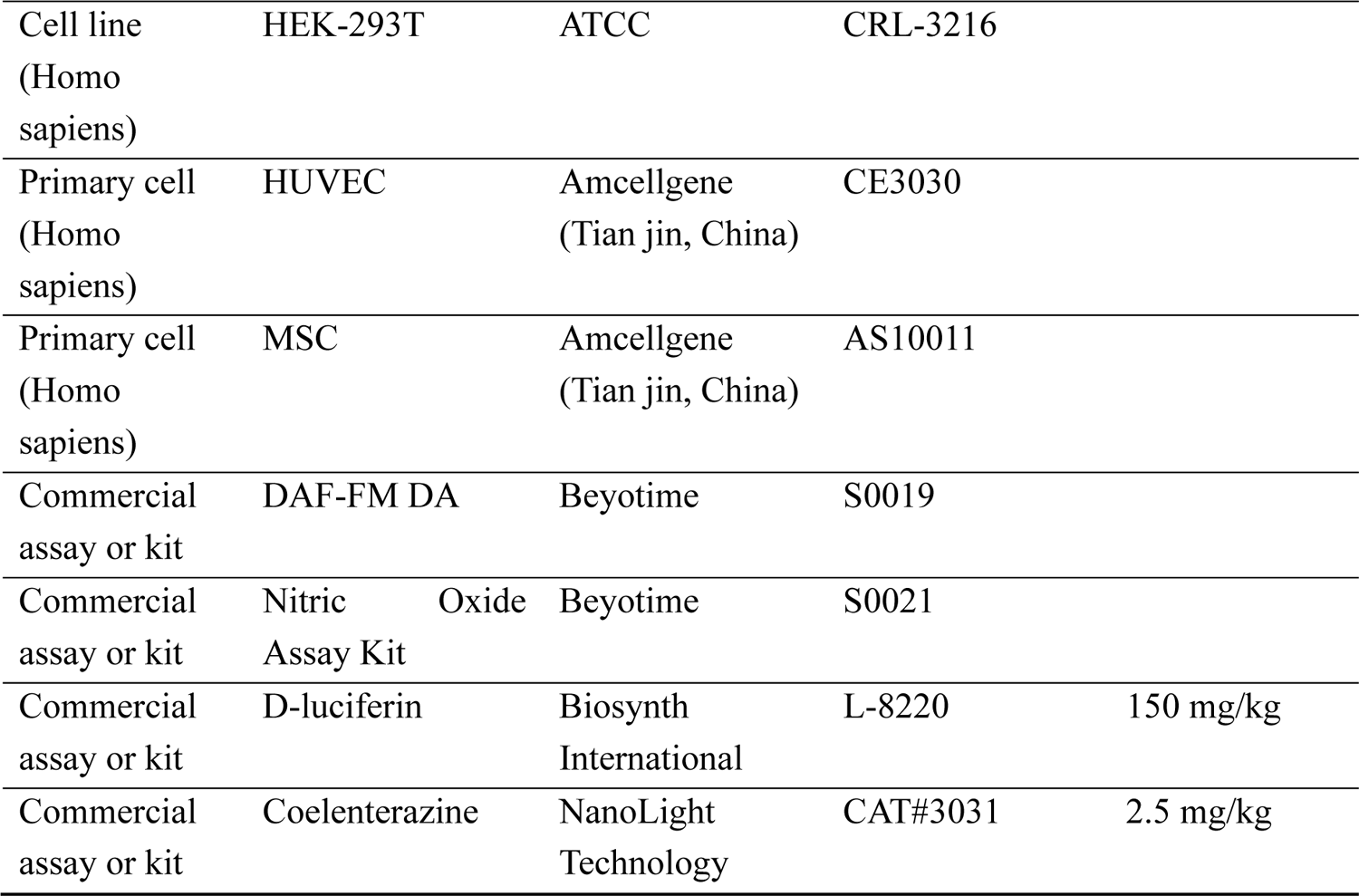

## Supplementary Information

**Supplemental Table 1.**
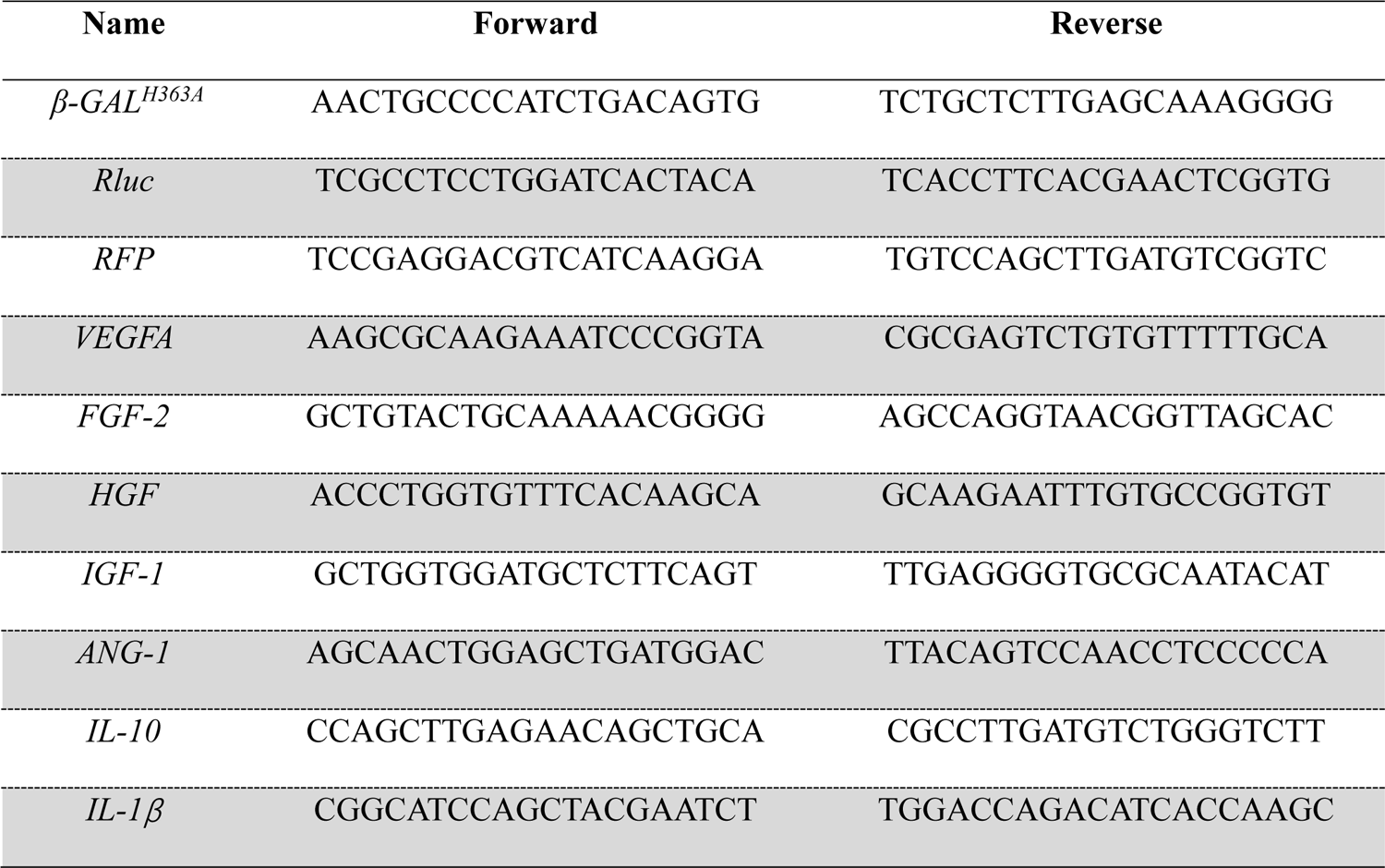
List of human primer sequences used for qPCR analysis in this study.

**Supplemental Table 2.**
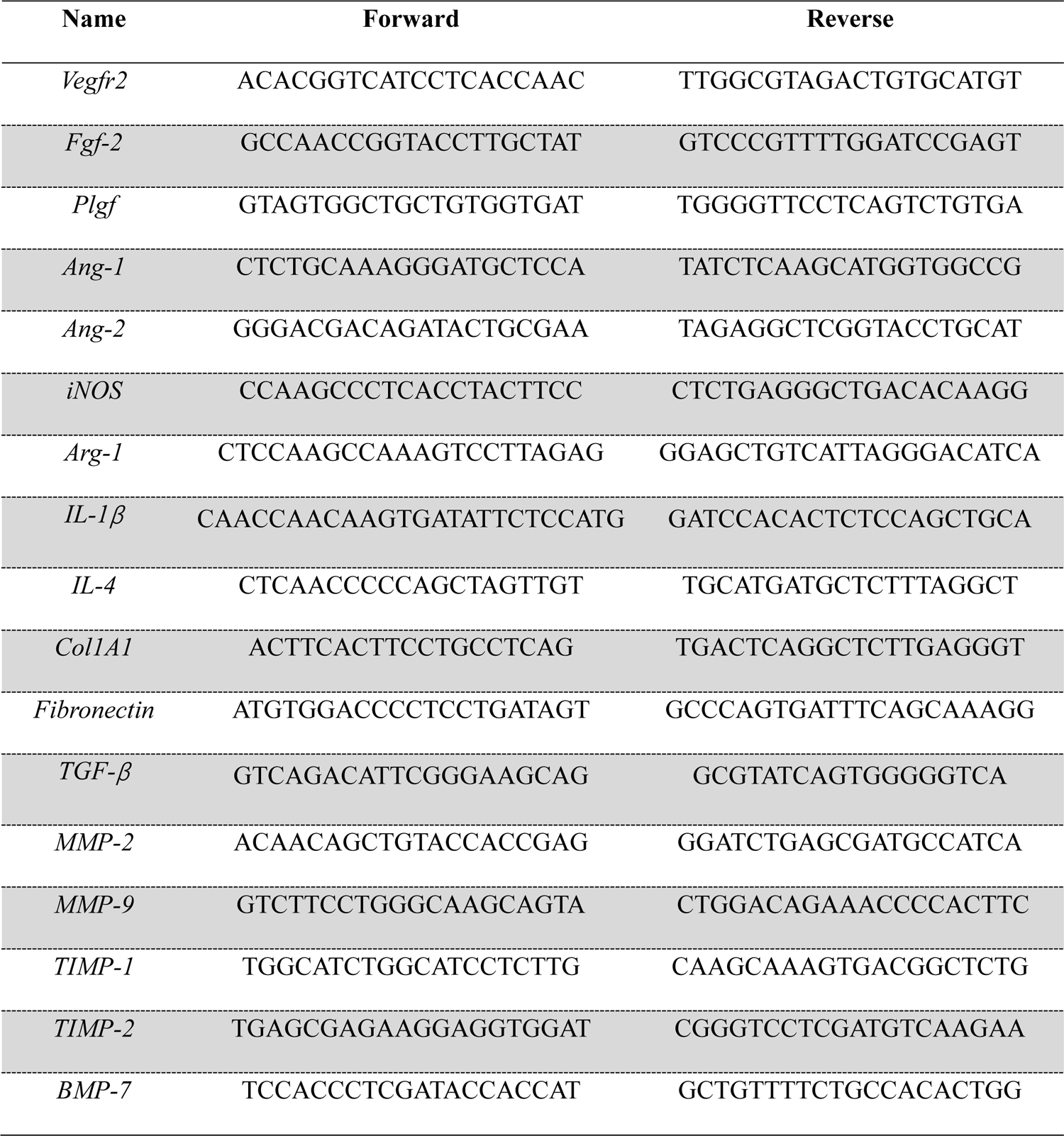
List of mouse primer sequences used for qPCR analysis in this study.

## Supplemental Methods

### Cell culture

#### HUVEC culture

Primary human umbilical vein endothelial cells (HUVECs) were purchased from the American Type Culture Collection (ATCC, Manassas, VA) and grown in EGM2 medium (Lonza, Walkersville, MD). Cells from passages 4 to 7 were used for the experiments in this study.

#### HK-2 culture

A human proximal tubule epithelial cell line (HK-2) was purchased from ATCC and cultured in DMEM/F12 medium (Gibco) supplemented with 10% FBS, penicillin (100 U/mL), and streptomycin (100 μg/mL).

#### HEK-293T culture

Human embryonic kidney 293T cells (HEK 293T) were obtained from ATCC and maintained in high-glucose DMEM with 10% FBS penicillin (100 U/mL) and streptomycin (100 μg/mL).

#### Construction of plasmids and engineered MSC (eMSC)

The coding sequence of mutant β-galactosidases (β-GAL^H363A^) can be obtained from previous publications(1). To achieve localization of the transplanted MSCs, we cloned the coding sequence of double-fusion reporter genes (Rluc-RFP: Renilla luciferase and red fluorescent protein) from a previously reported pcDNA3.1-RFP-Rluc-HSV-ttk plasmid (2, 3). Subsequently, the sequences encoding β-GAL^H363A^ and Rluc-RFP were inserted into the lentiviral backbone plasmid, which was carried out by Miaolingbio Company (Wuhan, China). The lentiviral vector was cotransfected with the second-generation lentivirus packing vectors psPAX2 and pVSVG into 293T cells by Lipofectamine 2000 (Invitrogen, Grand Island, NY) to produce lentivirus. The infection efficiency was evaluated by RFP expression, as directly observed under an inverted fluorescence microscope (Olympus, Japan), and further confirmed by β-GAL^H363A^ expression (C-His-tagged recombinant protein), as evidenced by immunofluorescence.

#### Gene expression analysis

Total RNA was isolated from kidney tissues or cells using TRIzol reagent (Invitrogen) according to the manufacturer’s instructions and spectrophotometrically quantified using a NanoDrop One (Thermo, USA), and RNA integrity was tested by 1% agarose gel electrophoresis. For RNA reverse transcription into complementary DNA (cDNA), 1.0 μg of total RNA was treated with the BioScript All-in-One cDNA Synthesis SuperMix kit (Bimake, Houston, TX). Real-time (RT)-quantitative polymerase chain reaction (qPCR) analysis was performed using a CFX96TM Real-Time System (Bio-Rad, Hercules, USA) in a 20 μL reaction volume containing qPCR SYBR green master mix (Yeasen, Shanghai, China). The 2^-ΔΔCt^ method was used to determine the relative mRNA folding changes. Primers are listed in **Supplementary Tables 1 and 2**.

#### Western blotting analysis

MSCs were collected and homogenized in RIPA lysis buffer containing protease inhibitors (Solarbio, Shanghai, China), and a BCA Protein Assay Kit (Thermo Scientific) was used to quantify the total protein. The total proteins were diluted in 4× SDS‒PAGE loading buffer, boiled for 10 minutes, electrophoresed on 10% polyacrylamide gels and blotted on 0.2 μm polyvinylidene fluoride (PVDF) membranes (Millipore, Darmstadt, Germany). After blocking with 5% nonfat milk for 2 h at room temperature, blots were incubated with primary antibodies overnight at 4 °C followed by secondary antibodies at room temperature for 2 h. Anti-6×His tag (β-GAL^H363A^) antibody was used. Signals were generated by using enhanced chemiluminescence (ECL) reagent (Millipore, USA) and were captured by using a Tanon-5200 Chemiluminescence Imaging System (Tanon Science & Technology Co Ltd., Shanghai, China). Quantification was performed using ImageJ software.

#### Renal oxidative stress-related index

Kidney samples (100 mg) were collected and homogenized in 900 μL cold PBS with a tissue homogenizer to prepare 10% tissue homogenate. Then, the activities of the oxidative stress-related factors MDA, SOD, and GSH were detected with a lipid peroxidation MDA Assay Kit (A003-1-2, Nanjing Jiancheng Bioengineering Institute), SOD Assay Kit (A001-3-2, Nanjing Jiancheng Bioengineering Institute), and GSH Assay Kit (A005-1-2, Nanjing Jiancheng Bioengineering Institute).

#### *In vitro* proangiogenic assay

To analyze the pro-angiogenic effect of eMSCs *in vitro*, a scratch wound-healing assay was performed to evaluate the migration capacity of HUVECs using a coculture system. HUVECs were seeded in a 6-well plate cocultured with eMSCs or eMSCs with MGP administration on a 24-mm Transwell (3412, Corning). When HUVECs reached confluence, scratch wounds were generated using the tip of a 10 μL micropipette. Images of 5 microscopic fields per well were taken at 0 h and 12 h, and the percentage of wound healing was quantitated using ImageJ software.

For the tube formation assay, 150 μL/well growth factor-reduced cold Matrigel (Corning, Corning, NY) was coated onto 48-well plates and then incubated at 37 °C for 30 min for gelatinization. Then, 3 × 10^4^ HUVECs were seeded per well and cocultured with eMSCs or eMSCs with MGP administration in a 6.5-mm Transwell (3413, Corning). After incubation for 12 h, the number of branches and nodes was measured in three randomly selected microscopic fields using ImageJ software as described before.

#### Histopathology and Immunostaining

Mice were euthanized to harvest the tissue samples at the indicated time points. For immunofluorescent staining, 5 μm cryosections were washed three times in PBS. Next, incubated with 5% BSA containing 0.1% Triton X-100 at RT for 1h, and then incubated with a series of primary antibodies overnight at 4 °C. The following secondary antibodies used were Alexa Fluor 488, Alexa Fluor 594, Alexa Fluor 555, and Alexa Fluor 647 labeled goat anti-rabbit, goat anti-mouse, or goat anti-rat (1:400; Invitrogen). The FITC-labeled Lotus tetragonolobus lectin (LTL, 1:400, Vector Laboratories, Burlingame, CA) was used to stain proximal tubules. DAPI was used for nuclear counterstaining. For quantification, the positive immunostaining area or positively stained cells were measured by ImageJ software.

For immunohistochemistry staining kidney tissues, after dewaxing and rehydration, paraffin sections were incubated with primary antibodies overnight at 4 °C. Horseradish peroxidase (HRP)–conjugated secondary antibody was used (1:200, SP-9000, ZSGB-BIO, Beijing, China), and DAB staining kit was performed to develop colors (ZLI-9017, ZSGB-BIO). The integrated optical density (IOD) values of positive staining were measured using Image-pro plus 6.0 software.

#### Annexin V apoptosis analysis

For apoptosis analysis, 1 × 10^4^ eMSC cells were cultured on 12-well plates and then treated with 200 μM hydrogen peroxide (H_2_O_2_) for 12 h. Cells were collected and then costained with annexin V– fluorescein isothiocyanate (Annexin V)-FITC (CA1020, Solarbio) and 7-aminoactinomycin D (7-AAD) (ST515, Beyotime) for 30 min, and cell apoptosis rates were measured using a FACSCalibur flow cytometer (BD Biosciences) and analyzed using FlowJo software (Tree Star).

## SUPPLEMENTAL RESULTS

### Alleviated renal fibrosis

In the I/R injured kidney, deteriorated renal function and damaged renal architecture lead to tubulointerstitial inflammation, which further results in kidney fibrosis and the progression of chronic kidney disease (CKD). Masson trichrome staining demonstrated that interstitial fibrosis developed in the later stage after injury (28 d), while eMSCs with MGP administration markedly attenuated the area of renal fibrosis (**Fig. S9A**). The kidney sections stained with collagen type IV further confirmed the inhibition of renal fibrosis in the eMSCs with MGP administration group, similar to Masson trichrome staining (**Fig. S9B**). Furthermore, the mRNA expression of ECM synthesis and fibrosis pathway-related genes further confirmed the histologic pattern. The gene expression levels of collagen type 1-A1 (Col1A1), fibronectin, TGFβ, MMP-2 and MMP-9 in kidneys were enhanced in the PBS group but notably inhibited by NO-eMSC therapy (**Fig. S9C**). Meanwhile, the administration of eMSCs significantly increased the expression of antifibrotic genes, including BMP-7, TIMP-1, and TIMP-2, suggesting that eMSCs with MGP administration exerted superior antifibrotic actions in AKI (**Fig. S9D**). Conclusively, eMSCs with MGP administration therapy resulted in superior outcomes in promoting kidney regeneration via amelioration of renal function and suppression of renal fibrosis.

### Hemocompatibility and cytotoxicity of the NO prodrug, MGP

The hemocompatibility and cytotoxicity of the NO prodrug, MGP, were assessed before use in experiments. Hemolytic tests indicated that after incubation with Control and MPG, the cell integrity (**Fig. S14A**) and morphology (**Fig. S14B**) of red blood cells (RBCs) were similar to those of RBCs treated with negative control-PBS. As shown in **Fig. S14C**, there was no obvious change in the indexes of blood chemistry, including ALT and AST, for liver function in all groups, confirming the excellent biosafety of the NO-eMSC system. No histological abnormalities were observed in the main organs of any group (**Fig. S14D**), indicating that the eMSCs exhibited excellent blood compatibility.

## Supplementary Figure & Legends

**Fig. S1.**
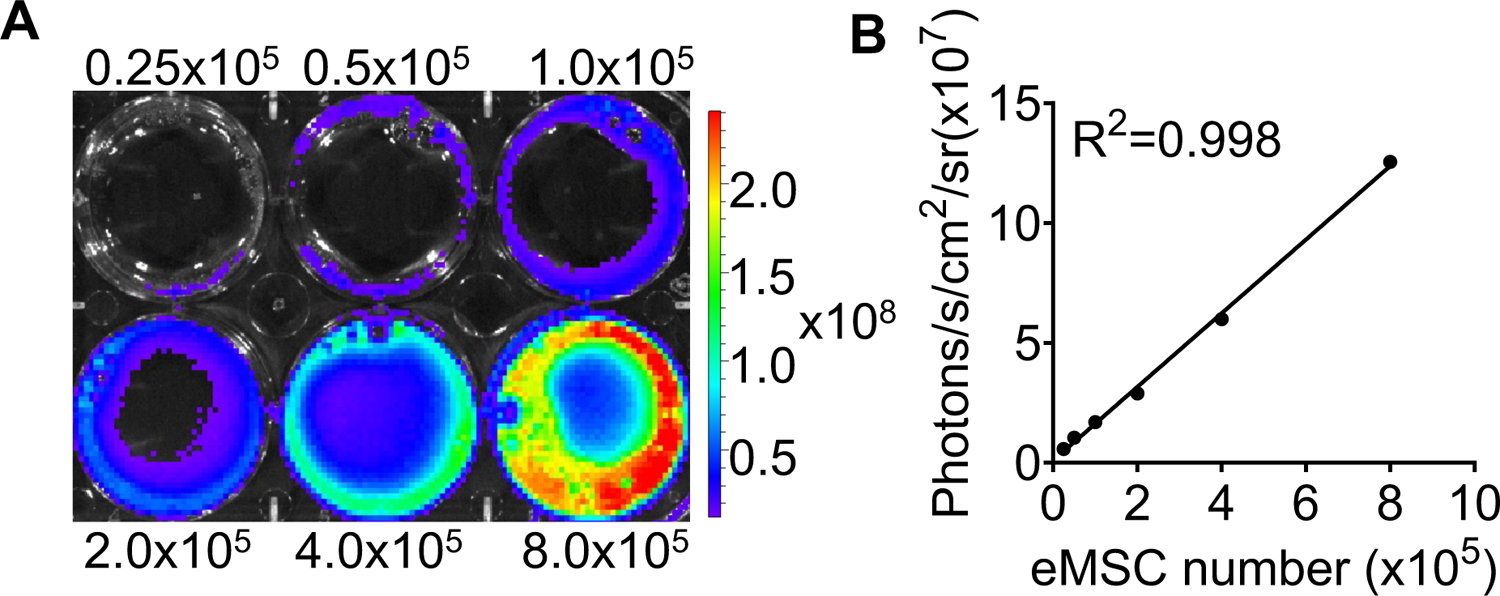
(**A** and **B**) *Ex vivo* bioluminescence imaging (BLI) of eMSCs showed a robust linear relationship (R^2^=0.998) with cell numbers in the presence of coelenterazine. All data are presented as the means ± SDs, \**P* < 0.05, n≥3 per group.

**Fig. S2.**
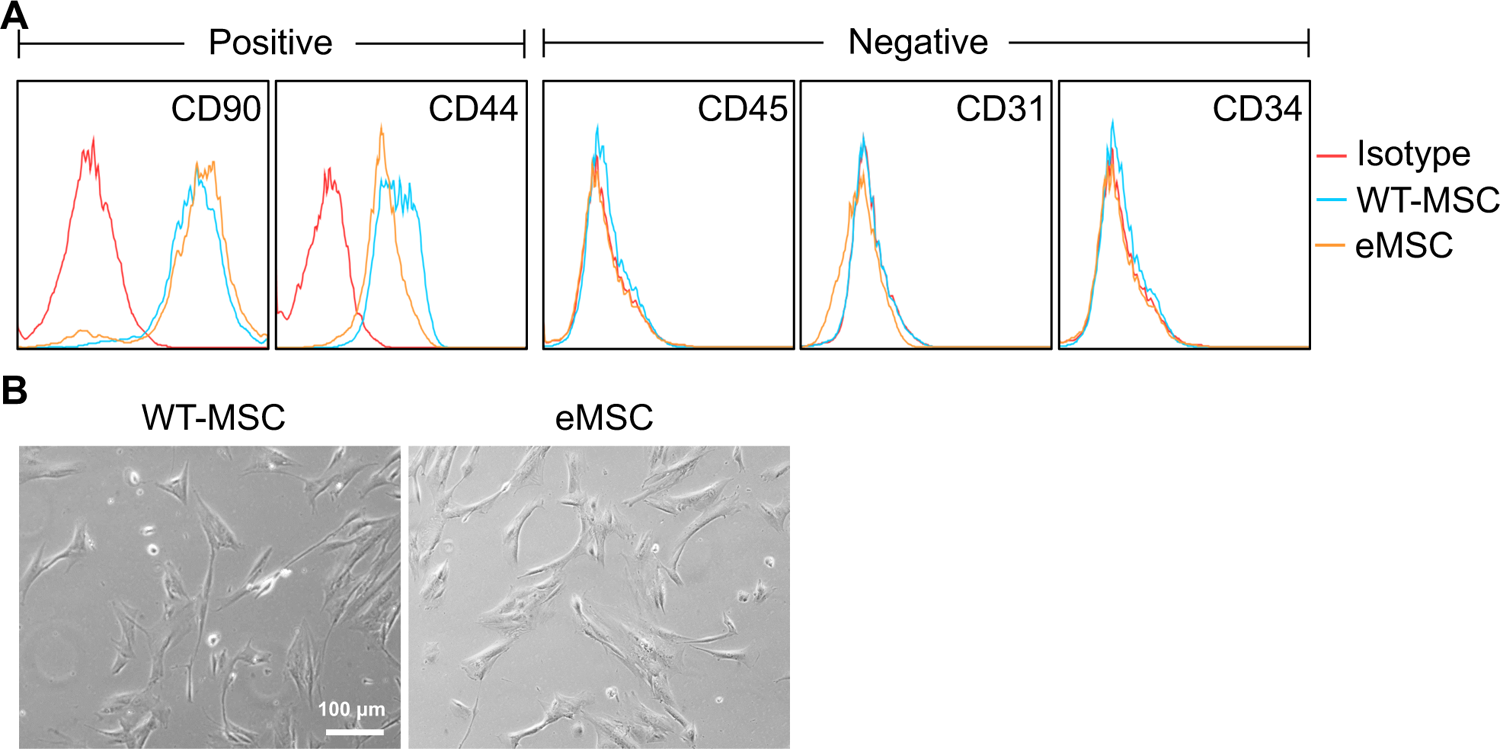
Characterization of the eMSC. (**A**) Comparison of immunophenotypes between wild type MSCs and eMSCs. Negative: CD45, CD31 and CD34; positive: CD90 and CD44; (**B**) Morphology of wild type MSCs and eMSCs. Scale bars, 100 μm. Data are expressed as the mean ± SD; data are not statistically significant when p values are not shown; n=3 per group.

**Fig. S3.**
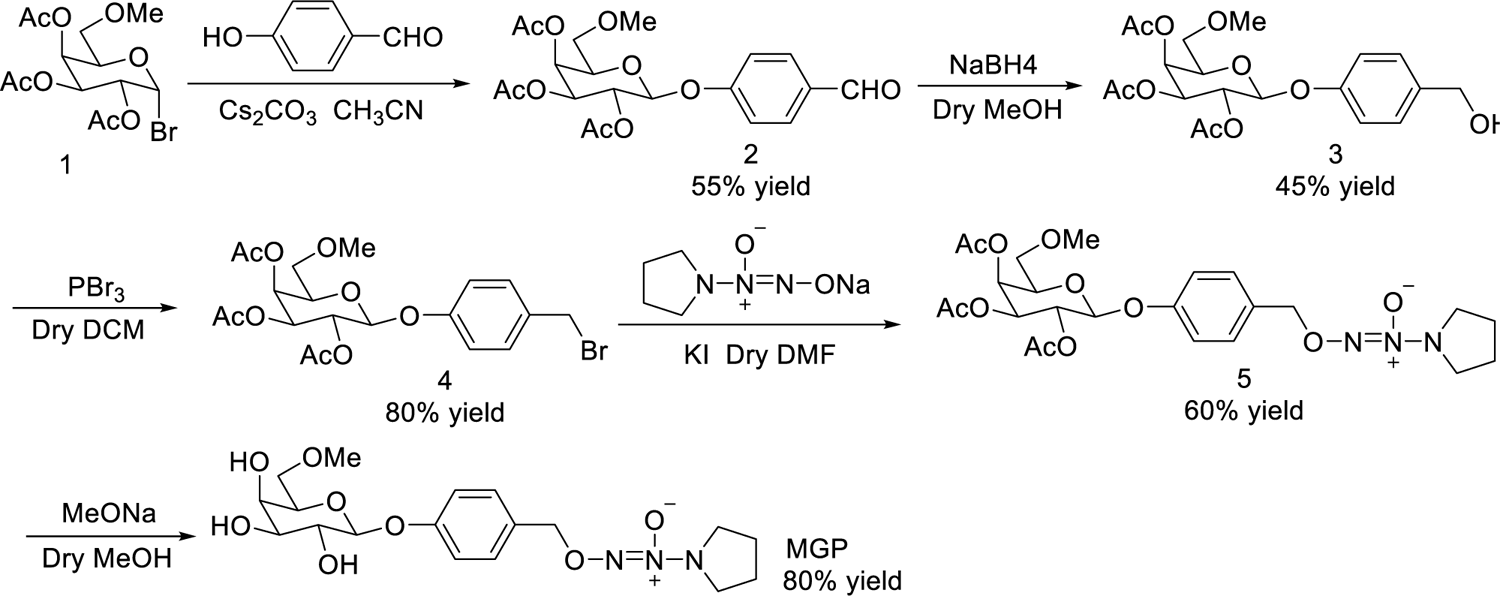
Synthesis route of the NO-prodrug MGP-NO.

**Fig. S4.**
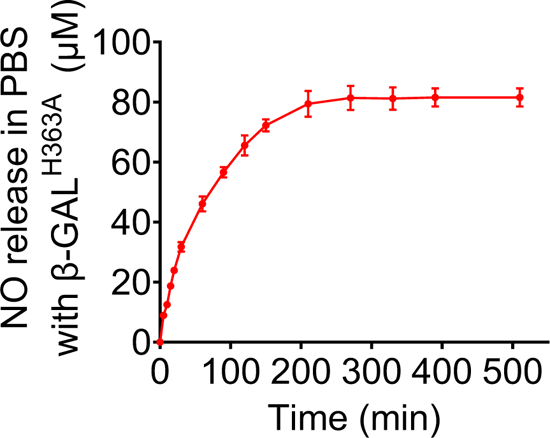
**(B)** The NO release profile of MGP-NO in response to β-GAL^H363A^ in PBS measured by Griess assay.

**Fig. S5.**
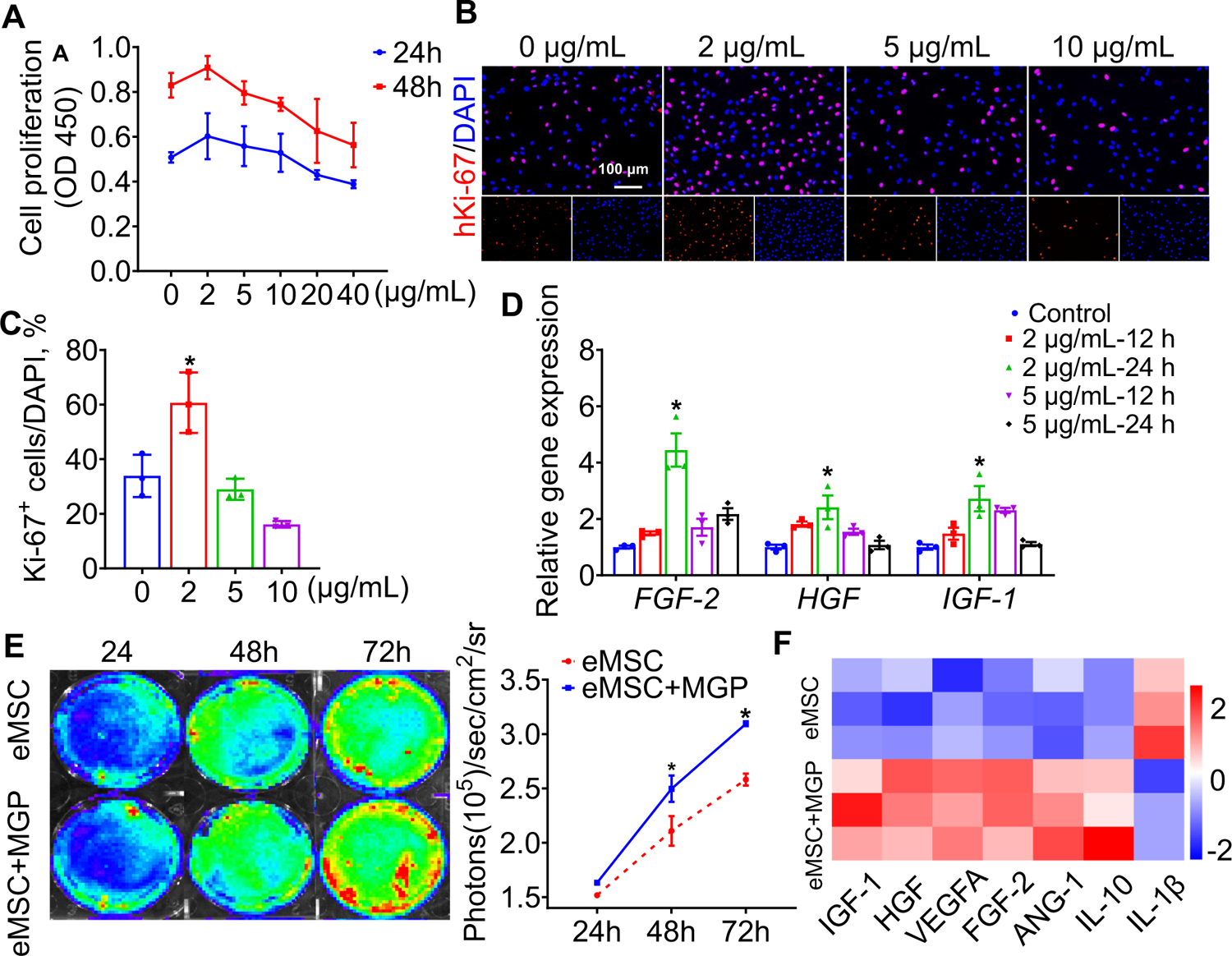
NO release facilitated the proliferation potential of eMSCs *in vitro*. (**A**) CCK-8 assay showed cell proliferation after administration of different concentrations of the NO-eMSC system. (B) Representative imaging of proliferation marker Ki-67 (red) immunofluorescence staining. Scale bars, 100 μm. (**C**) Quantification of Ki-67-positive cells. (**D**) Growth factor-related gene (bFGF, HGF, and IGF-1) mRNA levels in cells treated with different concentrations of the NO-eMSC system. (**E**) The proliferation of eMSCs and eMSCs with MGP administration (2 μg/mL) was analyzed by BLI (left); quantification of BLI signals (right); (**F**) real-time qPCR analysis of the expression of a panel of functional-related genes treated with eMSCs and eMSCs with MGP administration (2 μg/mL) for 24 h. All data are presented as the means ± SDs, \**P* < 0.05, n≥3 per group.

**Fig. S6.**
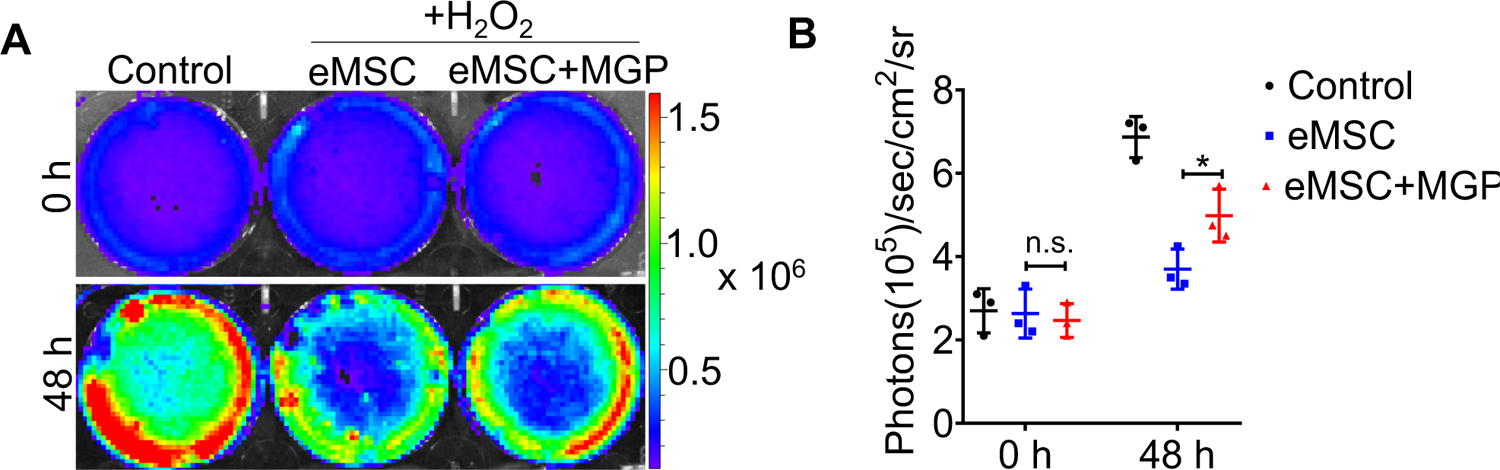
(**A** and **B**) The survival of eMSCs and eMSCs with MGP administration upon exposure to 200 μM H_2_O_2_ for 48 h was analyzed by BLI. All data are presented as the means ± SDs, \**P* < 0.05, n≥3 per group.

**Fig. S7.**
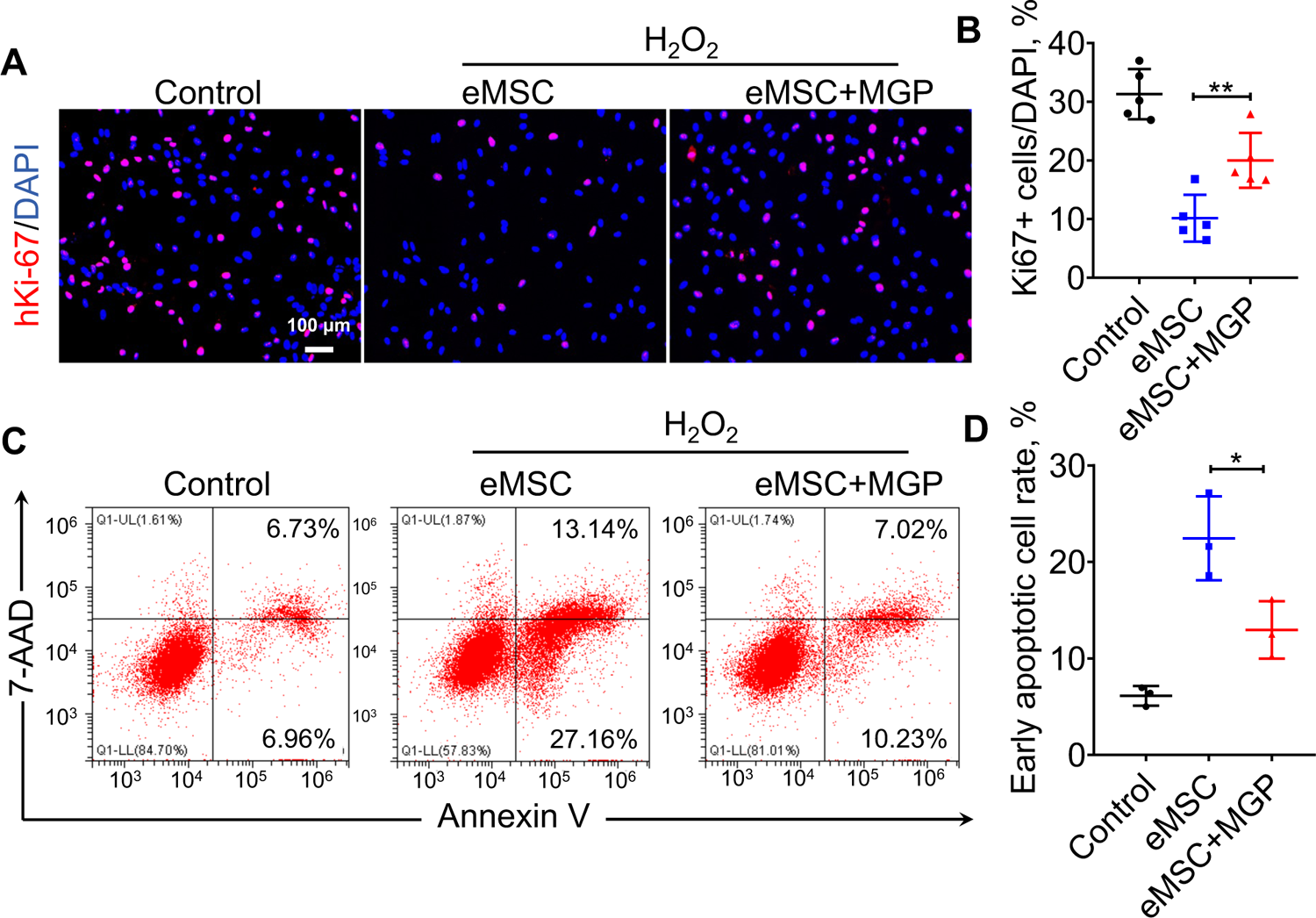
NO-eMSCs displayed higher resistance to apoptosis *in vitro*. Ki67 staining and percentage (**A**) of Ki67-positive cells (**B**) of eMSCs and eMSCs with MGP administration following exposure to 200 μM H_2_O_2_ for 24 h. Scale bars, 100 μm. (B) Annexin V/7-AAD double staining (**C**) and the percentages of apoptotic cells (**D**) upon exposure to 200 μM H_2_O_2_ for 12 h. All data are presented as the means ± SDs, n≥3 per group, \**P* < 0.05.

**Fig. S8.**
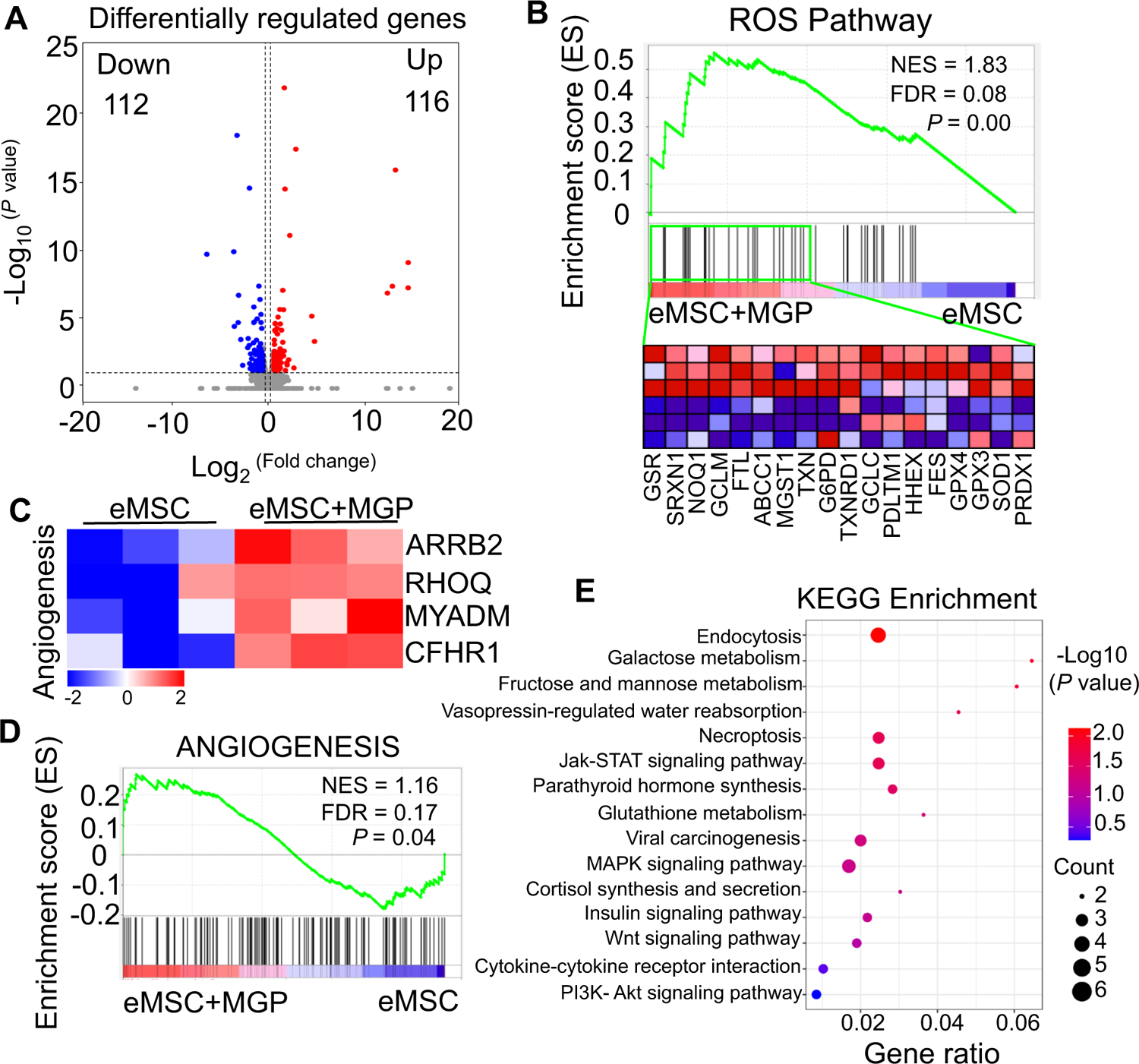
RNA-seq analysis revealed that the NO-eMSC system improved eMSC functionality. **(A)** Volcano plot illustrating differentially expressed gene (DEG) expression between eMSCs and eMSCs with MGP administration. **(B)** Gene set expression analysis (GSEA) revealed enrichment for the ROS pathway. FDR, false discovery rate; NES, normalized enrichment score. **(C)** Heatmap of representative angiogenesis-related genes. **(D)** GSEA revealed enrichment for angiogenesis pathways. FDR, false discovery rate; NES, normalized enrichment score. **(E)** Kyoto Encyclopedia of Genes and Genomes (KEGG) pathway annotation of DEGs.

**Fig. S9.**
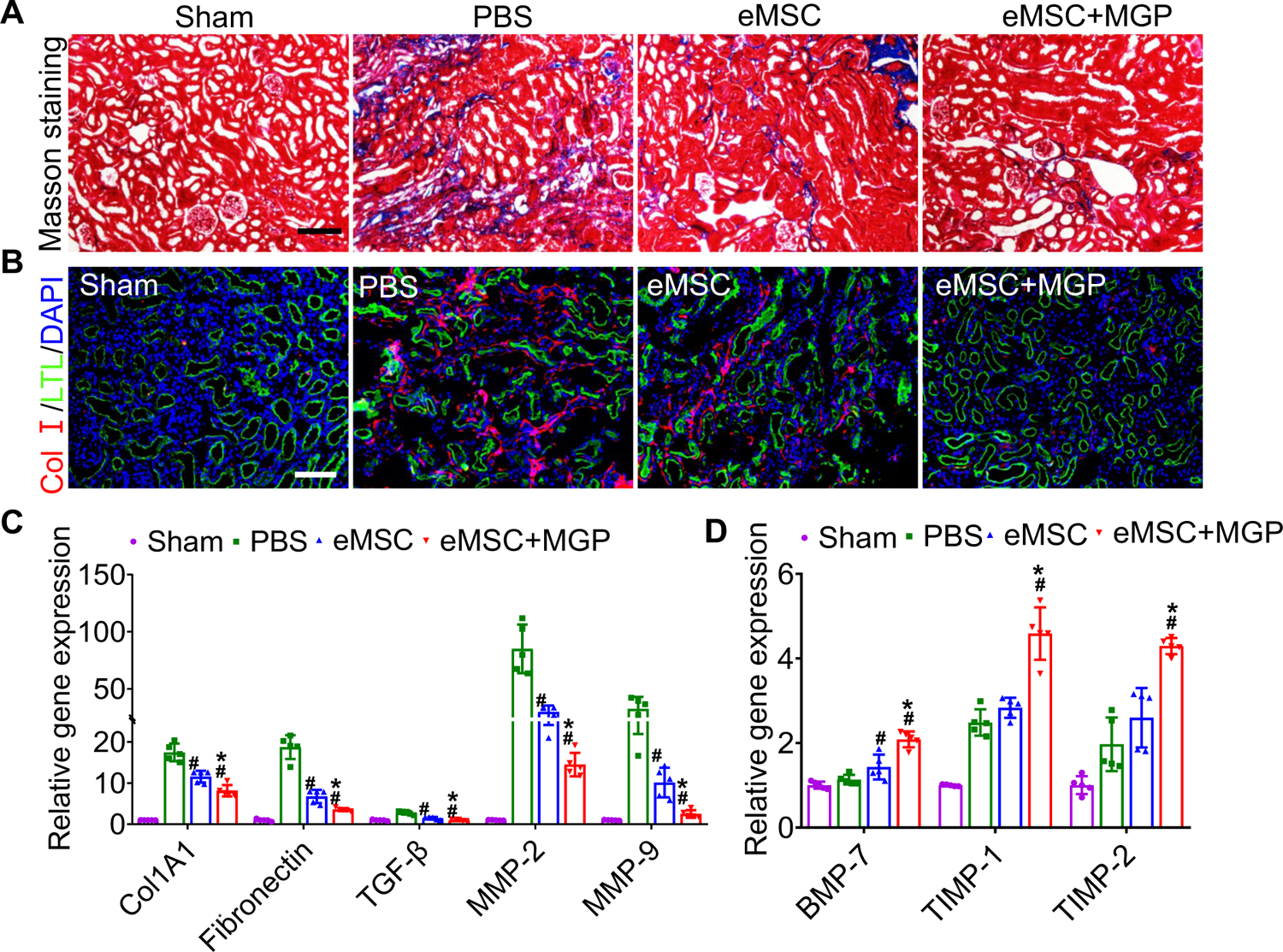
eMSCs with MGP administration attenuate renal fibrosis. (**A**) Masson trichrome staining was performed to examine kidney fibrosis on day 28 post AKI. Scale bar, 100 μm. (**B**) COL-I immunofluorescent staining was performed to examine kidney fibrosis on day 28 post AKI. Scale bar, 100 μm. (**C** and **D**) Real-time PCR analysis of fibrosis-related gene expression in the kidney at day 28 after AKI. All data are presented as the means ± SDs, n≥3 per group, \**P* < 0.05 versus eMSC; *^#^P* < 0.05 versus PBS.

**Fig. S10.**
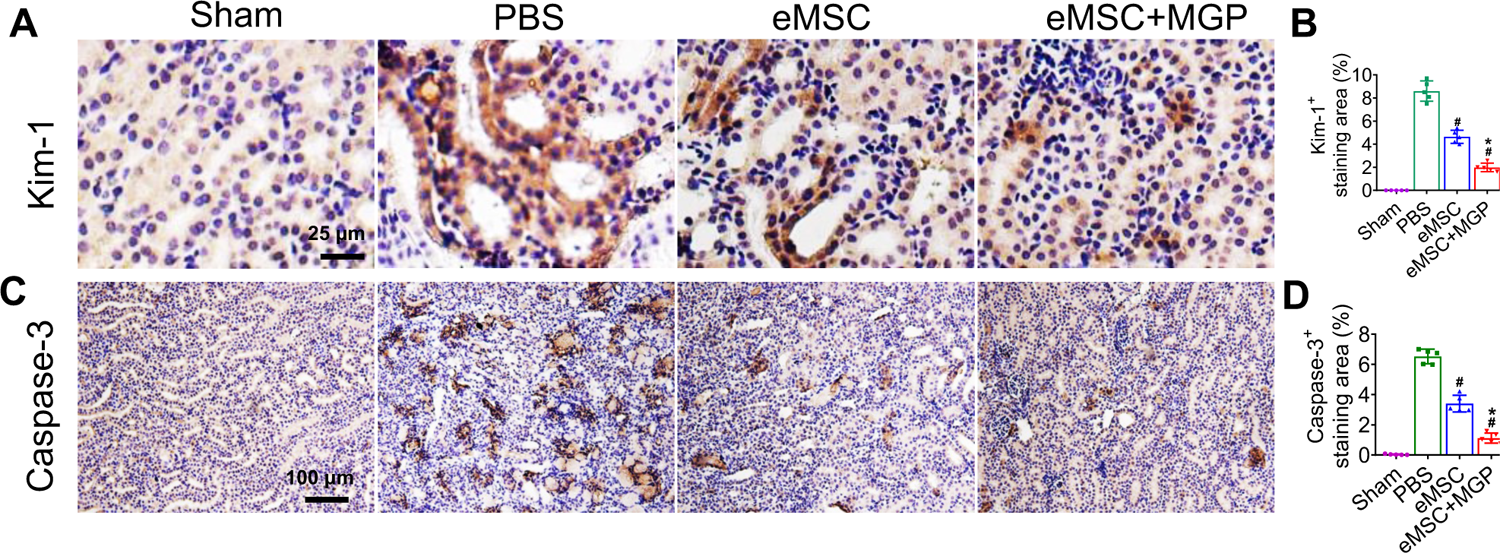
The NO-eMSC system protected against kidney injury by rescuing renal tubules. (**A, B**) Representative images of Kim-1 immunohistochemical staining on day 3 after AKI. Scale bar, 25 μm. (**C, D**) Caspase-3 immunohistochemical staining on day 3 after AKI. Scale bar, 100 μm. All data are presented as the means ± SDs, n≥3 per group, \**P* < 0.05 versus eMSC; *^#^P* < 0.05 versus PBS.

**Fig. S11.**
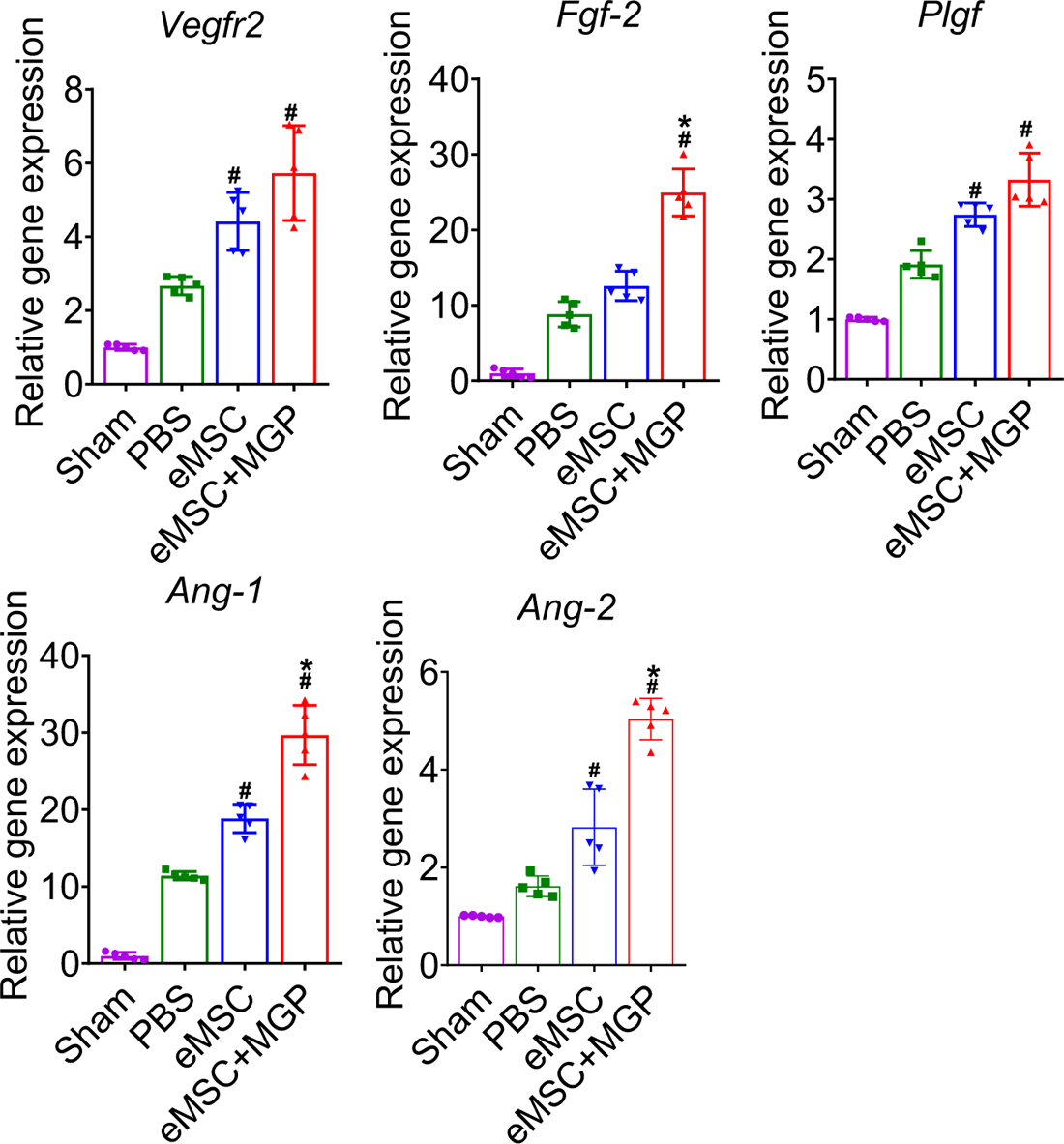
Enhanced angiogenic effects of the NO-eMSC system. mRNA levels of angiogenesis-related genes (VEGFR2, bFGF, PLGF, Ang-1, and Ang-1) in kidney tissues on day 7 after AKI. All data are presented as the means ± SDs, n=3 per group, \**P* < 0.05 versus eMSCs; *^#^P* < 0.05 versus PBS.

**Fig. S12.**
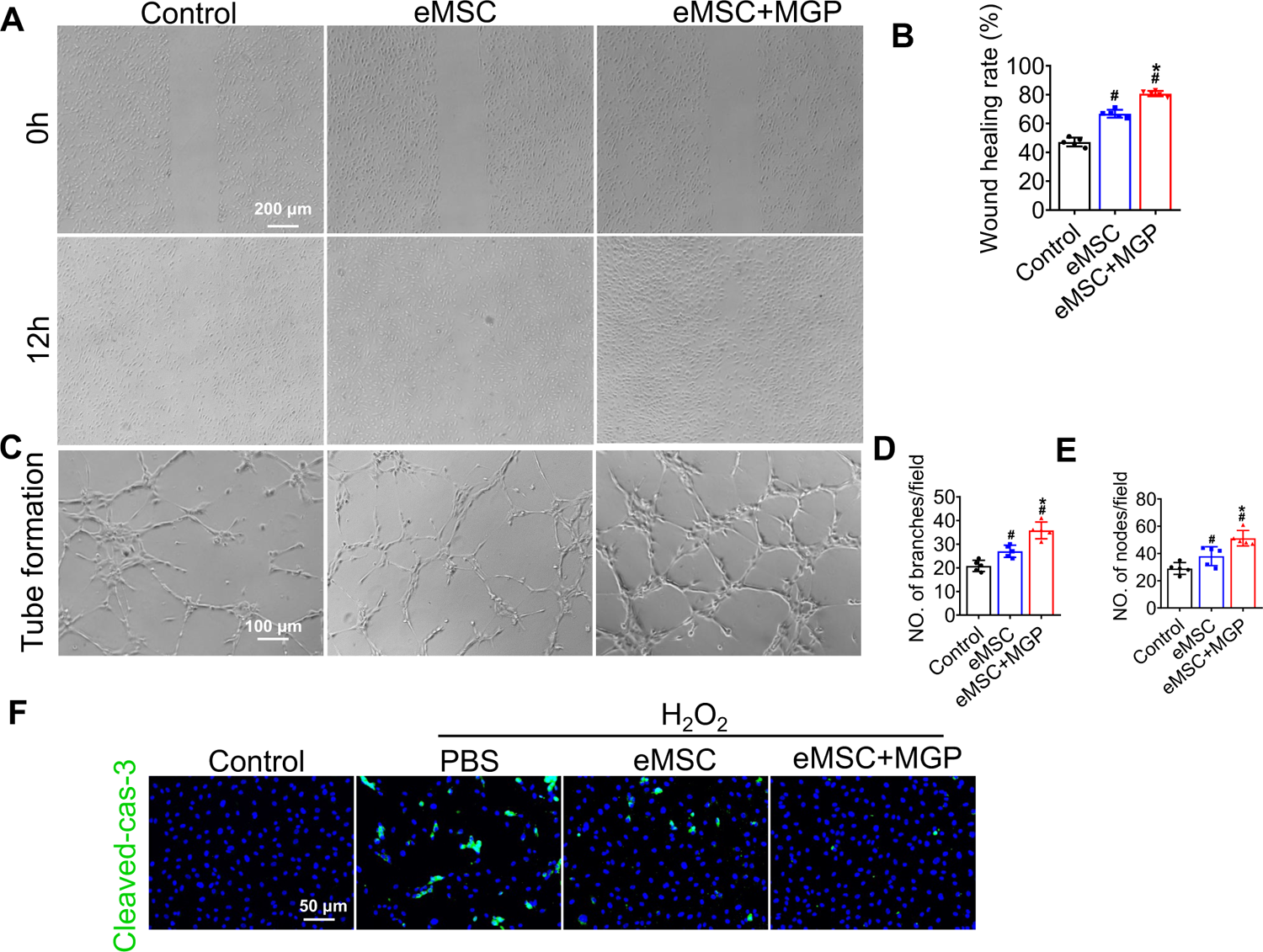
eMSCs with MGP administration enhanced the proangiogenic potential of HUVECs *in vitro*. (**A**) Representative images of the wound-healing migration assay in HUVECs cocultured with eMSCs and eMSCs with MGP administration. Scale bar, 200 μm. (**B**) Quantitative analysis of the wound-healing percentage ratio. (**C**) Representative images of tube formation of HUVECs cocultured with eMSCs and eMSCs with MGP administration. Scale bar, 100 μm. (**D** to **E**) Quantitative analysis of node number (D) and branch number (E) in three random fields in each group. (**F**) Cleaved caspase-3 immunostaining in HUVECs cocultured with eMSCs and eMSCs with MGP administration following exposure to 200 μM H_2_O_2_. Scale bars, 50 μm. DAPI demarcates nuclei. All data are presented as the means ± SDs, \**P* < 0.05 versus eMSC; ^#^*P* < 0.05 versus control. n≥3 per group.

**Fig. S13.**
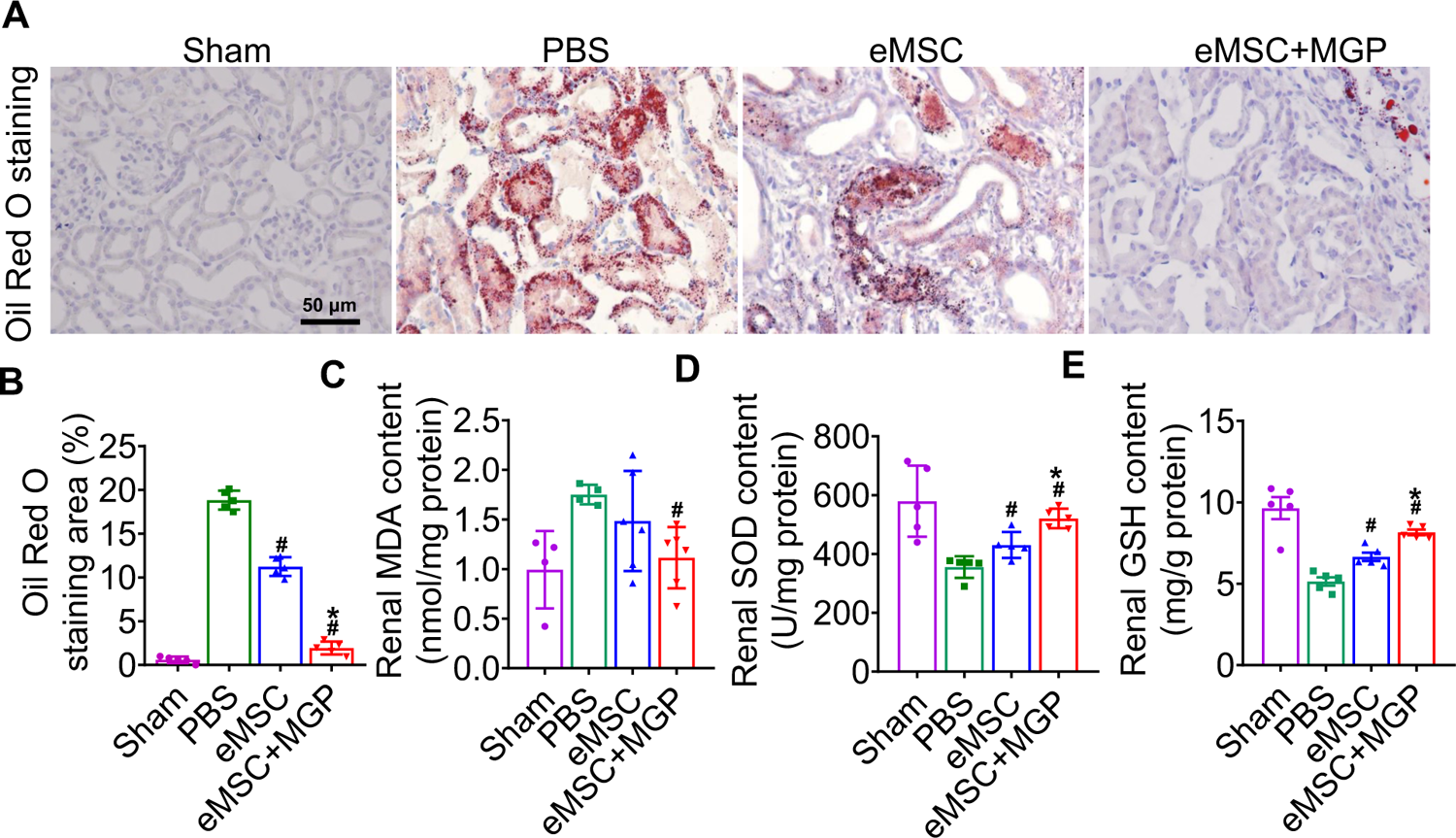
The NO-eMSC system ameliorates oxidative stress by protecting against antioxidation. (**A**) Representative images of Oil red O staining in proximal tubules at 3 days after AKI. Scale bars, 50 μm. (**B**) Quantification of the Oil Red O-positive area. Renal MDA (**C**), SOD (D) and GSH (**E**) content 3 days after AKI. All data are presented as the means ± SDs, n=3 per group, \**P* < 0.05 versus eMSCs; *^#^P* < 0.05 versus PBS.

**Fig. S14.**
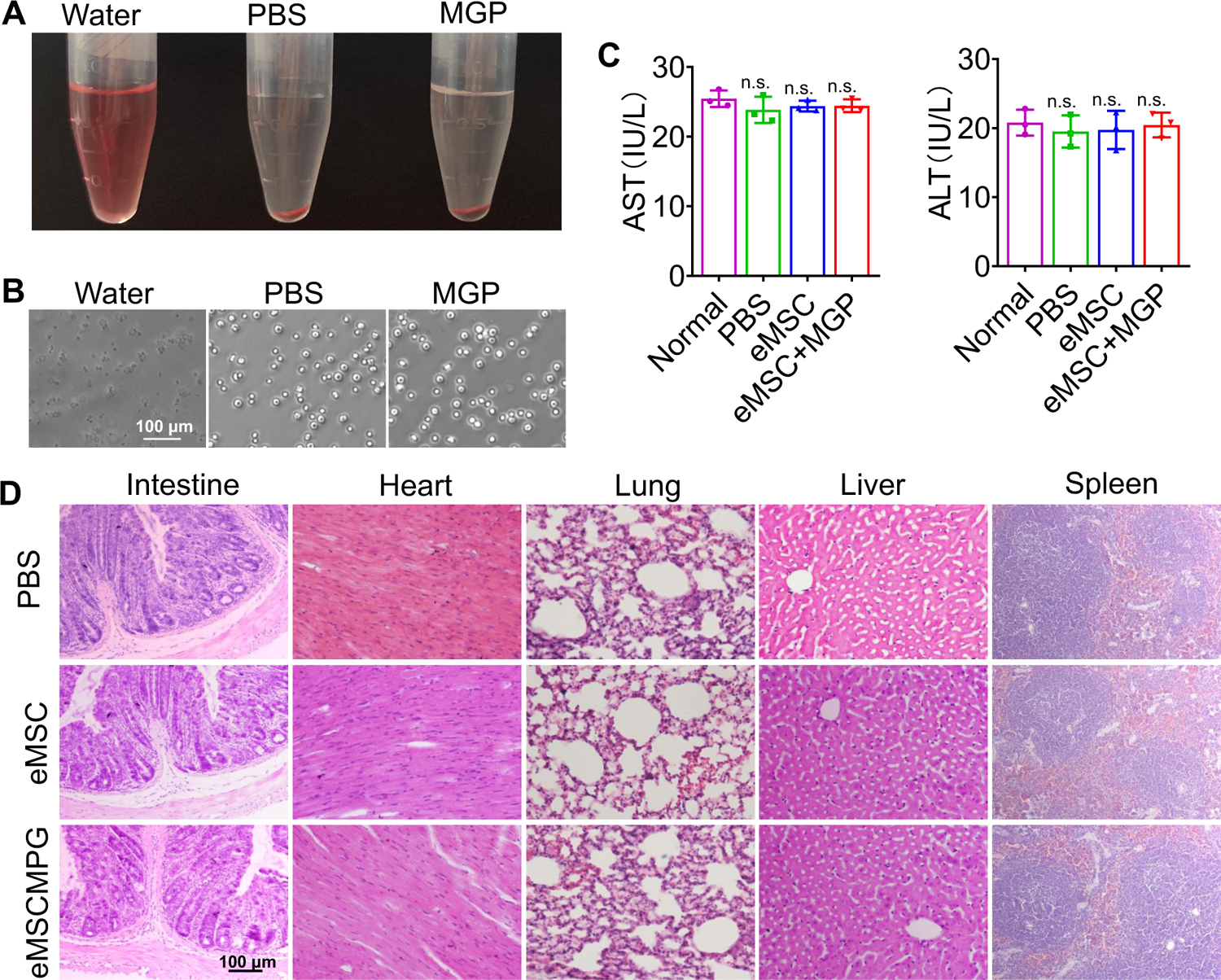
Hemocompatibility and cytocompatibility analyses of the NO-eMSC system. (**A**) Overview of the red blood cell hemolysis assay; red blood cells were incubated with water (positive control), PBS (negative control), and MGP (NO-prodrug). (**B**) Optical images of red blood cells. Scale bar, 100 μm. (**C**) Liver function indexes, including AST (left) and ALT (right) levels, were measured at day 28 after treatment. (**D**) Histopathological examination of the main organs (heart, liver, spleen, lung and intestine) to estimate the biosafety.

